## Supplemental Figures for "Widespread multi-targeted therapy resistance via drug-induced secretome fucosylation"

**Supplemental information**

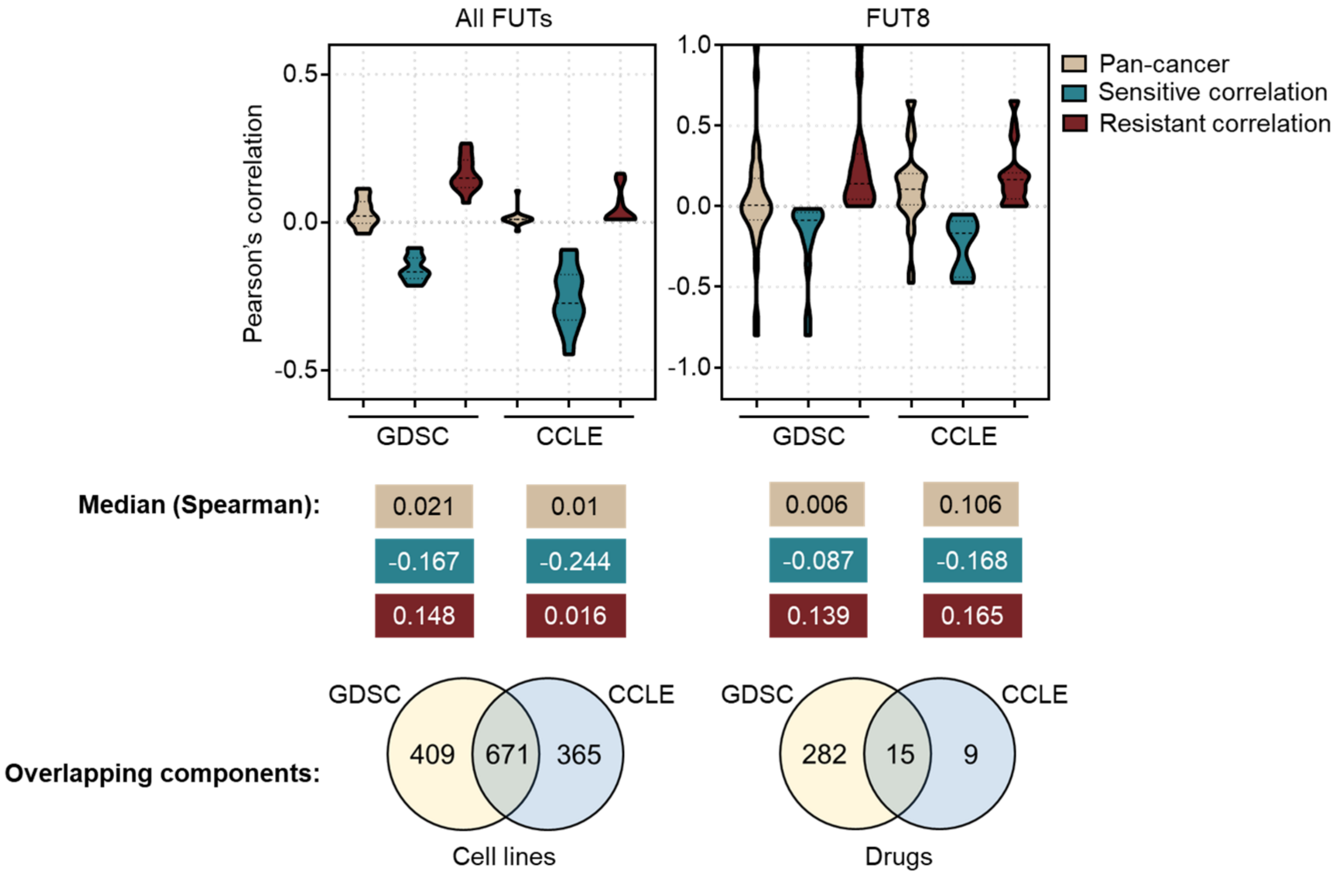


**Supplemental Fig. 1.** Comparison of correlations between FUT expression profiles and drug sensitivity and other shared components in GDSC and CCLE datasets. Top panel shows Pearson's correlation distributions and median values for comparable associations between FUT gene expression and drug sensitivity. Bottom panel shows cell line and drug overlaps between data sources.

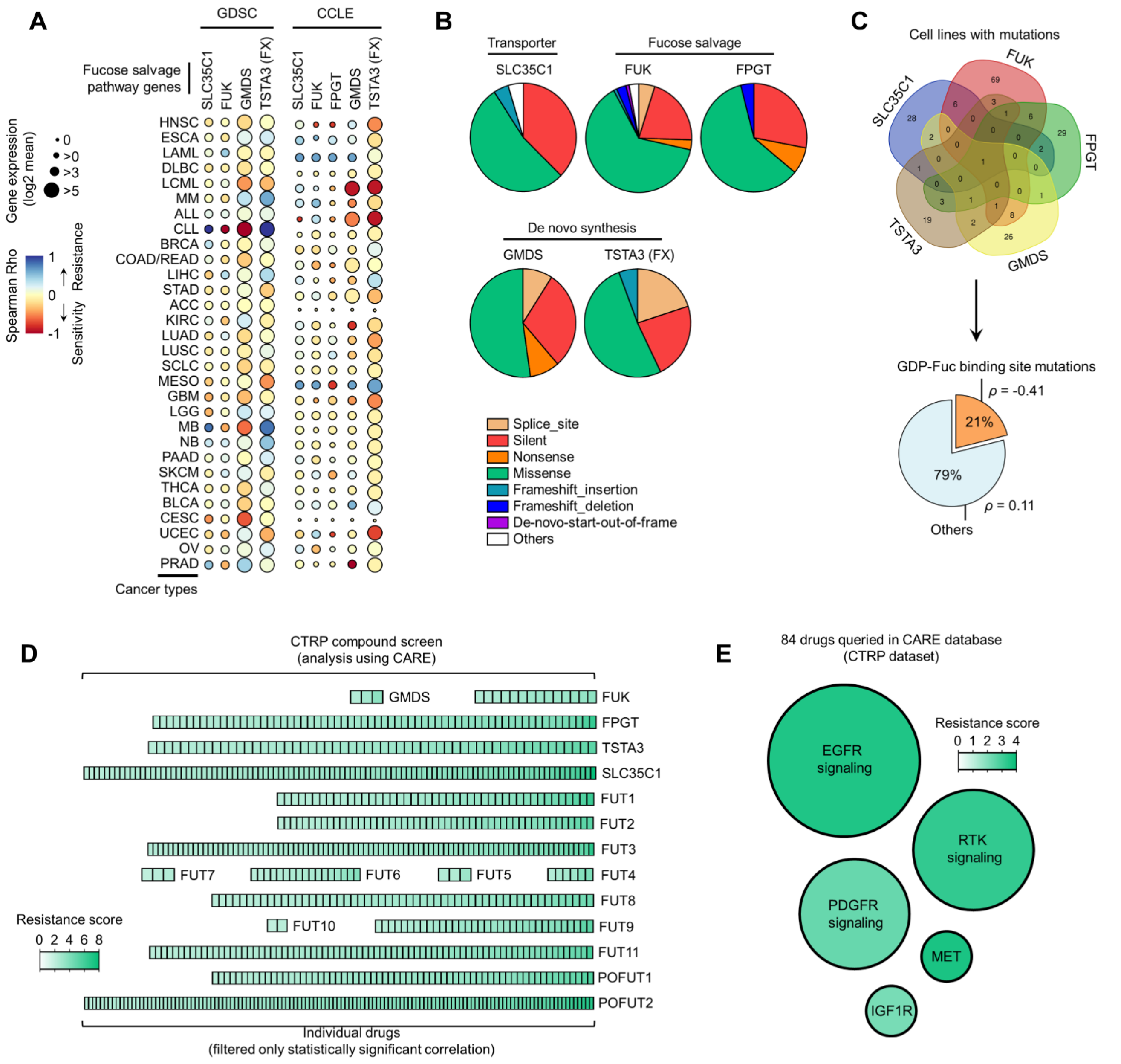


**Supplemental Fig. 2.** Correlation between fucose salvage and de novo synthesis gene expression and drug sensitivity.

(**A**) Heat-scatterplot visualization of correlation between indicated fucosylation gene expression and drug response per cancer type screened in GDSC and CCLE. Per-sample estimates of area under the fitted dose response curve were used as metric of drug response per cell line. Size of circle refers to mean log2 gene expression while color corresponds to Spearman's rank coefficients. Only statistically significant correlations are shown ($P<0.05$).

(**B**) Relative mean proportion of mutational signatures of indicated fucosylation genes per cancer type queried in CCLE.

(**C**) FUT mutations were classified as "GDP-Fuc binding site mutations" if any mutations (amino acid change) occurred near (±5 amino acid position) or at the annotated GDP-Fuc binding sites. Domain information was queried in UniProt. Spearman's rank coefficients (correlation between fucosylation gene expression and drug response) were calculated in cell lines carrying these mutations as opposed to those that do not ("others").

(**D**) Heatmap visualization of CARE scores showing correlation between indicated fucosylation gene expression and drug response for an individual targeted therapy in all cancer types screened in CTRP. Only statistically significant correlations are shown ($P<0.05$).

(**E**) Heat-scatterplot visualization of mean CARE scores of fucosylation genes as in D and drugs categorized according to their target signaling in all cancer types screened in CTRP. Only statistically significant correlations are shown ($P<0.05$).

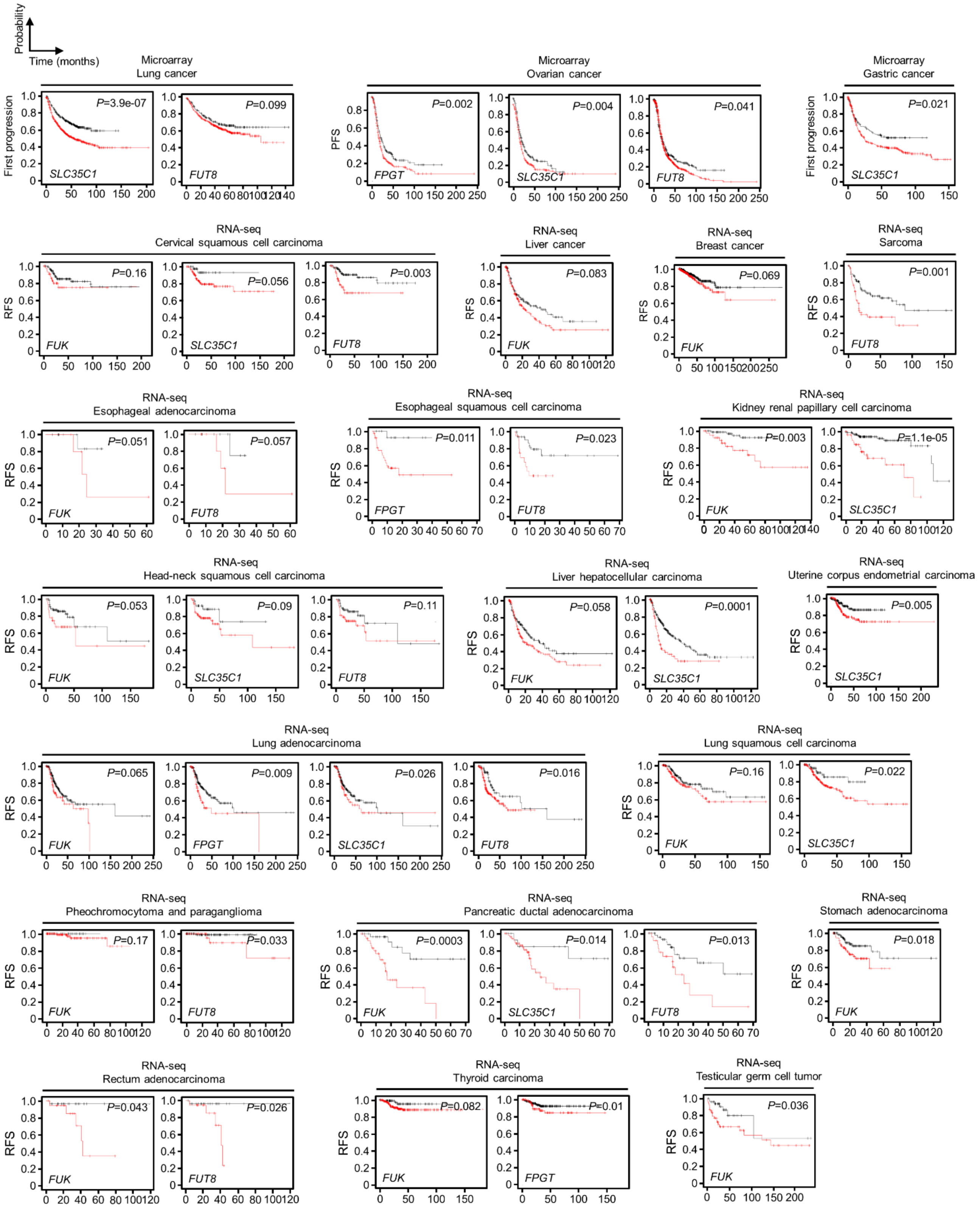


**Supplemental Fig. 3.** Fucose salvage gene expression is associated with cancer patient relapse and poor survival during/after therapy. Kaplan-Meier plots of first progression survival and RFS of patients diagnosed with indicated cancer types, stratified by the expression of indicated fucosylation gene in their primary tumors. *P*-values were calculated using a log rank test.

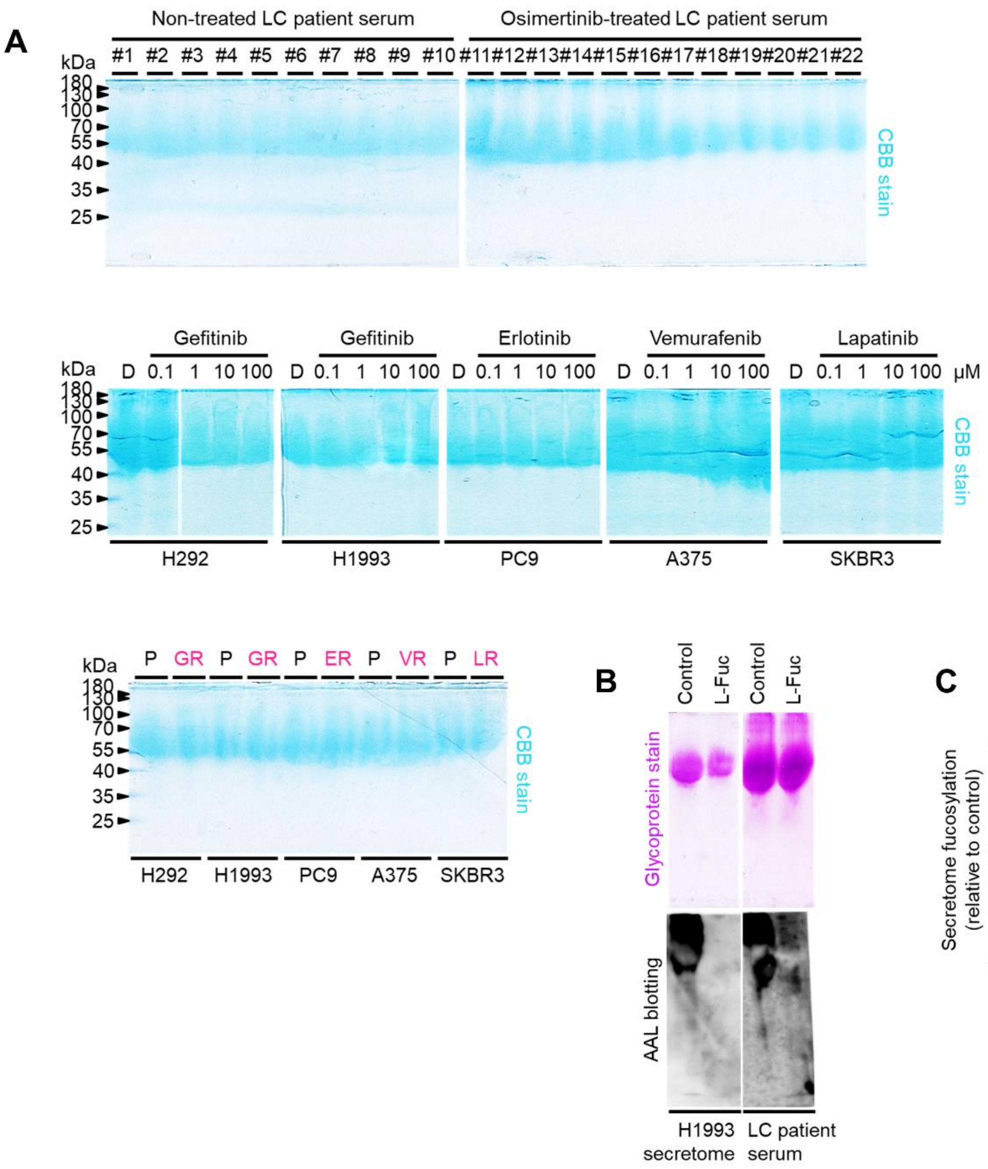


**Supplemental Fig. 4.** Equal loading controls and AAL specificity.
(**A**) Representative Coomassie stained SDS-PAGE gels showing fucosylated secretome proteins from indicated sensitive cells or DR clones prepared as in Fig. 1H following treatment with or without indicated drugs for 48 h.
(**B** and **C**) Characterization of AAL specificity by glycoprotein staining, AAL blot analysis, and sandwich ELLA. Cell secretomes were prepared as in Fig. 1H while patient sera were prepared as in Fig. 1F, except a competitive sugar (0.1 M L-Fuc) was added onto 15 μg N-glycoprotein samples prior to loading. Gels and blots are representative of two independent experiments. Values in sandwich ELLA are relative to control (means ± SD of two biological replicates). For statistical analysis, two-tailed Mann–Whitney *U* test was used.

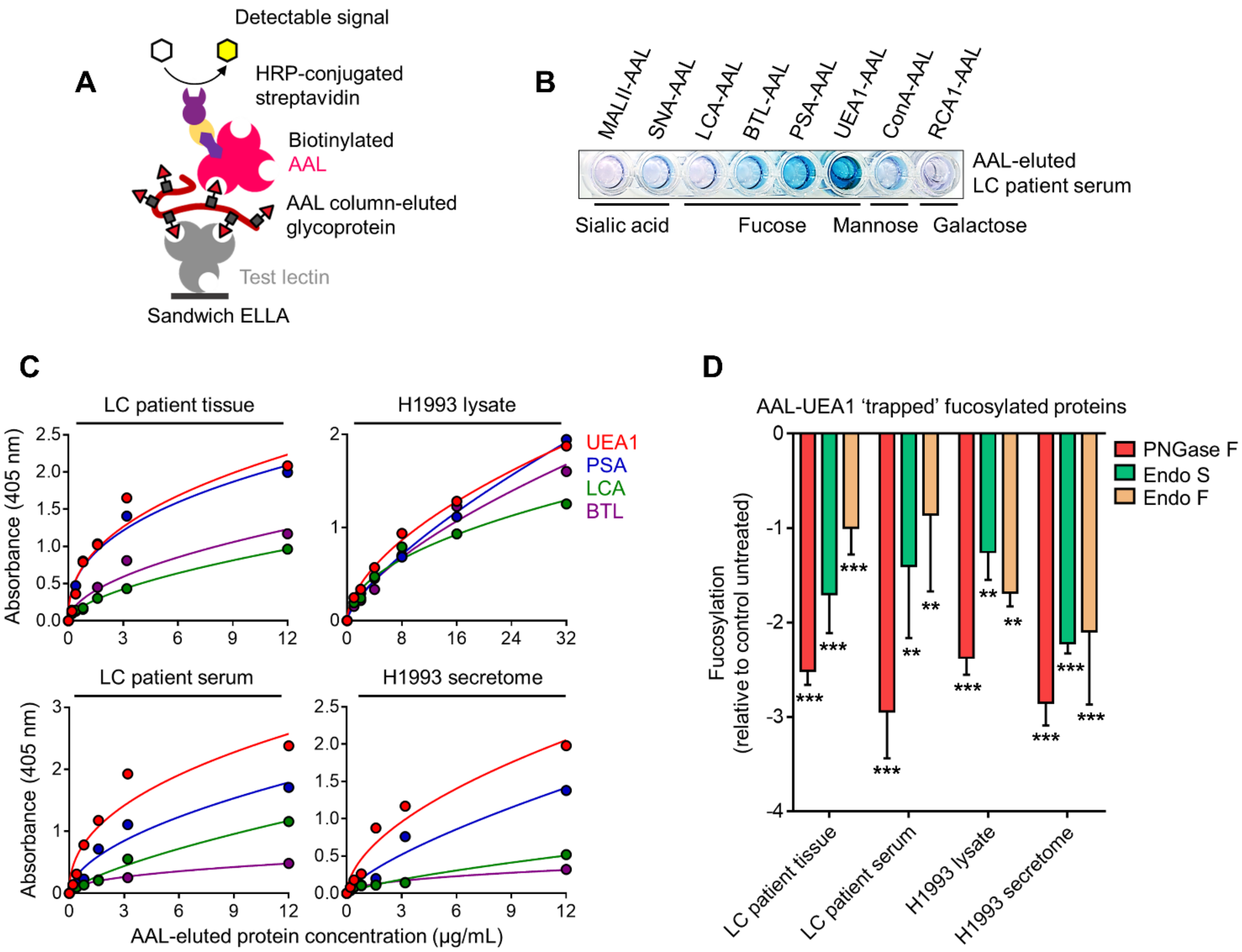


**Supplemental Fig. 5.** Sandwich ELLA.

(**A**) Schematic of sandwich ELLA.

(**B**) Representative detection of indicated captured glycoproteins from AAL-eluted LC patient serum (2 μg/mL per well) in a plate coated with indicated unconjugated test lectins.

(**C**) Characterization of fucosylation by sandwich ELLA in indicated AAL-eluted cell or tissue lysates, cell secretomes or patient sera. Values indicate mean absorbance at 405 nm from three replicates. Representative of two independent experiments.

(**D**) Characterization of fucosylation by sandwich ELLA in indicated AAL-eluted samples as in C upon exogenous de-N-glycosylation by (total 8U PNGase F or total 10U Endo S/F). Values are relative to untreated control (means ± SD of three biological replicates). ***P*<0.01, ****P*<0.001, Student's *t*-test.

**A**

| Type | Drug sensitive clone | Oncogenic background | Targeted therapy | Parental sensitivity | Drug resistant clone | Stepwise selection | Maintenance concentration | Xenograft validation | 3D tumorsphere validation |
|---|---|---|---|---|---|---|---|---|---|
| NSCLC | H292 | *WT* EGFR, *WT* KRAS | Gefitinib | Intermediate | H292-GR | >52 weeks | 2 μM | Yes[ref. 73] | Yes[this study] |
| NSCLC | H1993 | *WT* EGFR, high MET | Gefitinib | Intermediate | H1993-GR | >45 weeks | 2 μM | Yes[ref. 73] | Yes[this study] |
| NSCLC | HCC827 | *mut* EGFR | Gefitinib | Hyper | HCC8-GR | >41 weeks | 0.1 μM | Yes[ref. 74] | - |
| NSCLC | PC9 | *mut* EGFR | Erlotinib | Hyper | PC9-ER | >38 weeks | 0.1 μM | Yes[ref. 75] | Yes[this study] |
| NSCLC | H358 | *WT* EGFR, *mut* KRAS | Erlotinib | Intermediate | H358-ER | >27 weeks | 1 μM | - | - |
| NSCLC | H1975 | *mut* EGFR | Osimertinib | Hyper | H1975-OR | >40 weeks | 0.8 μM | - | - |
| NSCLC | HCC4006 | *mut* EGFR | Osimertinib | Hyper | HCC4-OR | >16 weeks | 1 μM | - | - |
| NSCLC | HCC827 | *mut* EGFR | Osimertinib | Hyper | HCC8-OR | >28 weeks | 1 μM | Yes[ref. 74] | - |
| NSCLC | H2009 | *mut* KRAS | Dasatinib | Intermediate | H2009-DsR | >12 weeks | 2 μM | - | - |
| NSCLC | H1993 | *WT* EGFR, high MET | PHA665752 | Hyper | H1993-PR | >18 weeks | 0.4 μM | - | - |
| NSCLC | EBC-1 | high MET | Crizotinib | Hyper | EBC1-CR | >21 weeks | 0.8 μM | - | - |
| NSCLC | H3122 | rearranged EML4-ALK | Crizotinib | Intermediate | H3122-CR | >17 weeks | 1 μM | - | - |
| NSCLC | HCC78 | rearranged SLC34A2-ROS1 | Crizotinib | Hyper | HCC7-CR | >17 weeks | 0.5 μM | - | - |
| Melanoma | A375 | *mut* BRAF | Vemurafenib | Hyper | A375-VR | >27 weeks | 1 μM | - | - |
| Melanoma | A375 | *mut* BRAF | Dabrafenib | Hyper | A375-DbR | >12 weeks | 1 μM | - | - |
| Breast adeno | SKBR3 | HER2+ | Lapatinib | Hyper | SKBR3-LR | >14 weeks | 0.7 μM | - | - |

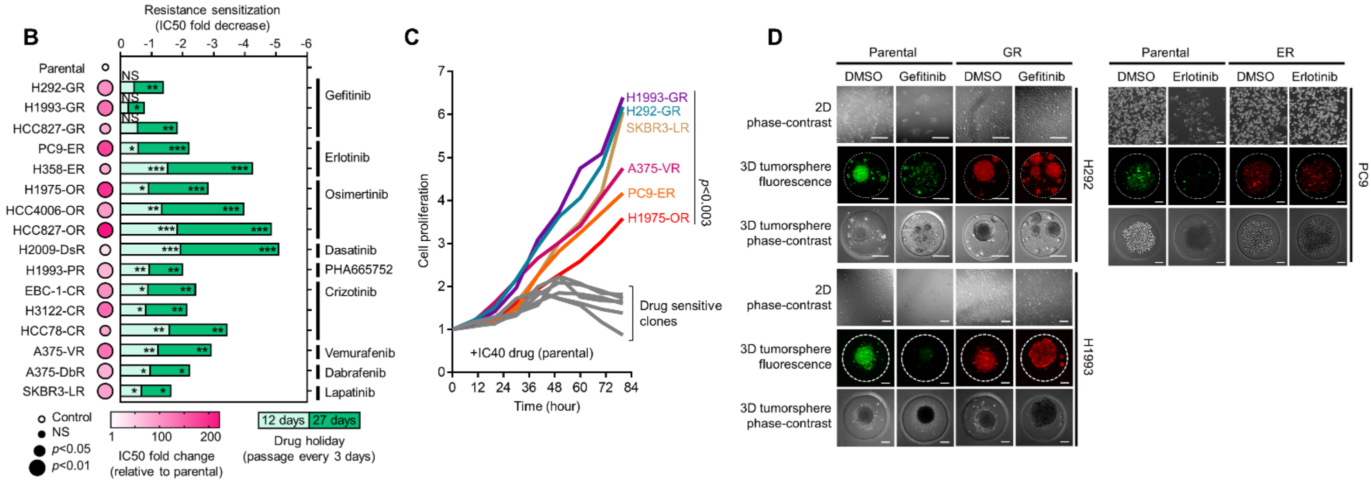


**Supplemental Fig. 6.** Characterization of DR clones.
(**A**) Summary of drug resistance selection in a panel of human cancer cell lines.
(**B**) Characterization of resistance in DR clones. IC50 values are relative to parental. *P* values are indicated as size of the corresponding circle; Student's *t*-test. NS, not significant. Stability of resistance was assayed using 12- and 27-day drug holidays. Cells were treated with or without drugs for 72 h with a concentration dilution series and were assayed by SRB. Representative of three independent experiments. *$P<0.05$, **$P<0.01$, ***$P<0.001$, Student's *t*-test.
(**C**) Proliferation of indicated sensitive cells or DR clones upon treatment with IC40 concentrations of indicated drugs (assessed in parental cells) for indicated times and were assayed by SRB. Values are relative to time point 0. Representative of two independent experiments. For statistical analysis, Student's *t*-test was used.
(**D**) Morphological evaluation of parental cells and GR clones in 2D and 3D cultures. For 3D spheroids, cells were transfected with CellTracker-Green for sensitive or CellTracker-Red for resistant. Cultures were treated with or without 2 μM gefitinib or 0.1 μM erlotinib for 72 h. Representative of two independent experiments.

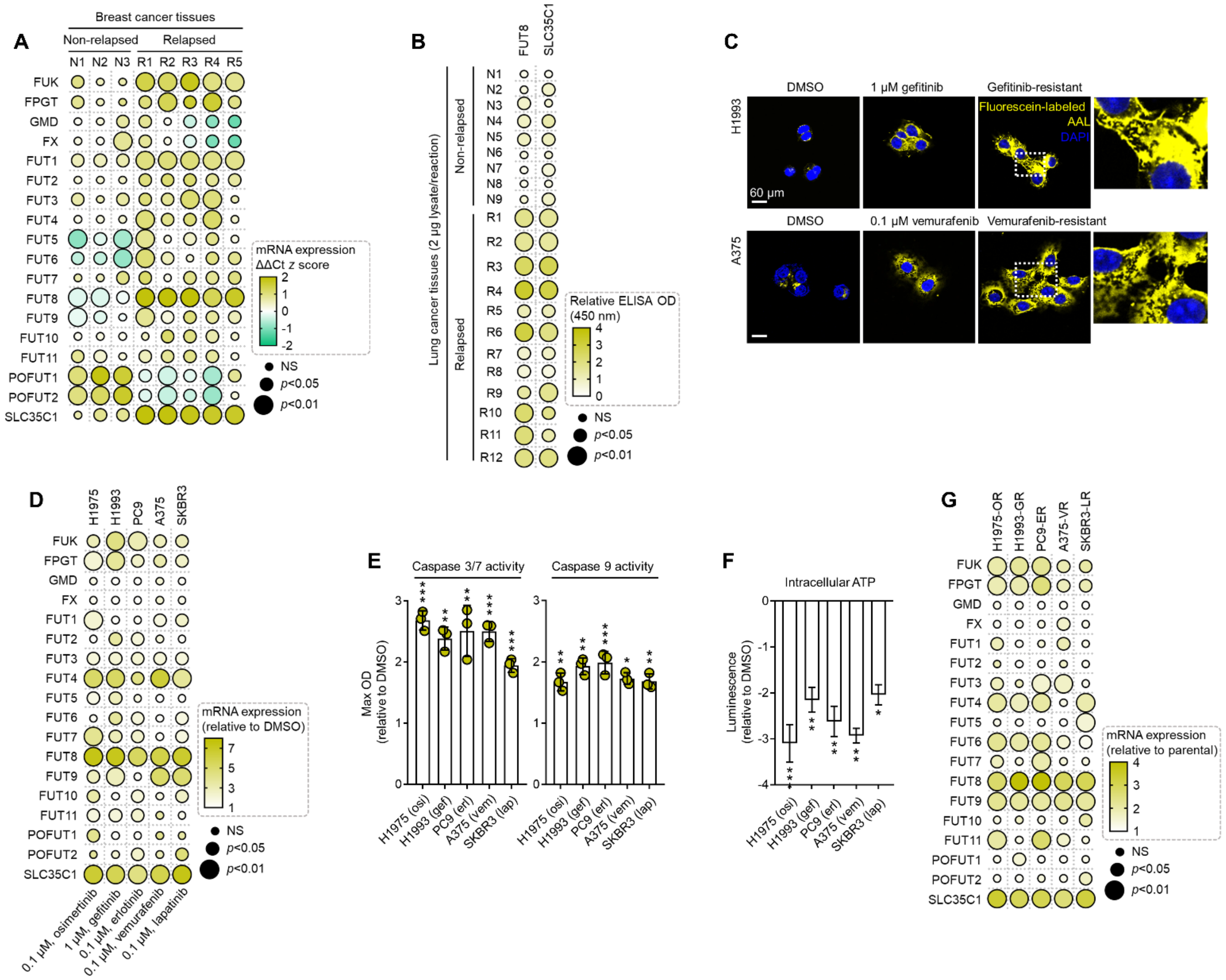

**Supplemental Fig. 7.** Expression and activity of fucose salvage genes and fucosyltransferases are associated with cancer patient relapse, therapy-induced apoptosis, and multiple acquired targeted therapy resistance.

(**A**) qPCR analysis of indicated fucosylation gene expression in indicated FFPE tumor tissue sections from patients with breast cancer who underwent sequential multidrug therapy. Log-transformed gene expression values are relative to the non-relapsed sample with lowest expression for the indicated gene (not displayed in the heatmap) and were normalized to GAPDH levels (means ± SD of three biological replicates). *P* values are indicated as size of the corresponding circle; Student's *t*-test. NS, not significant.

(**B**) ELISA analysis of FUT8 and SLC35C1 in indicated tumor tissues from patients with lung cancer who underwent first-line therapy. Values indicate mean absorbance at 450 nm from three replicates. Representative of two independent experiments. *P* values are indicated as size of the corresponding circle; Student's *t*-test. NS, not significant.

(**C**) Representative confocal images of indicated sensitive cells or DR clones stained for core fucosylation (fluorescein-conjugated AAL; yellow) and DAPI (nuclei; blue). Cells were treated with and without 1 μM gefitinib or 0.1 μM vemurafenib for 48 h. Representative of two independent experiments.

(**D**) qPCR analysis of indicated fucosylation gene expression in cells treated with or without indicated drug concentrations for 72 h. Values are relative to DMSO and were normalized to GAPDH levels (means ± SD of three biological replicates). *P* values are indicated as size of the corresponding circle; Student's *t*-test. NS, not significant.

(**E**) Caspase 3/7 DEVDase and caspase 9 activities of indicated cells and treatment conditions as in D. Values are relative to DMSO (means ± SD of two biological replicates). **P*<0.05, ***P*<0.01, ****P*<0.001, Student's *t*-test.

(**F**) Intracellular ATP activity of indicated cells and treatment conditions as in D. Values are relative to DMSO (means ± SD of two biological replicates). **P*<0.05, ***P*<0.01, ****P*<0.001, Student's *t*-test.

(**G**) qPCR analysis of indicated fucosylation gene expression in indicated DR clones. Values are relative to parental and were normalized to GAPDH levels (means ± SD of three biological replicates). *P* values are indicated as size of the corresponding circle; Student's *t*-test. NS, not significant.

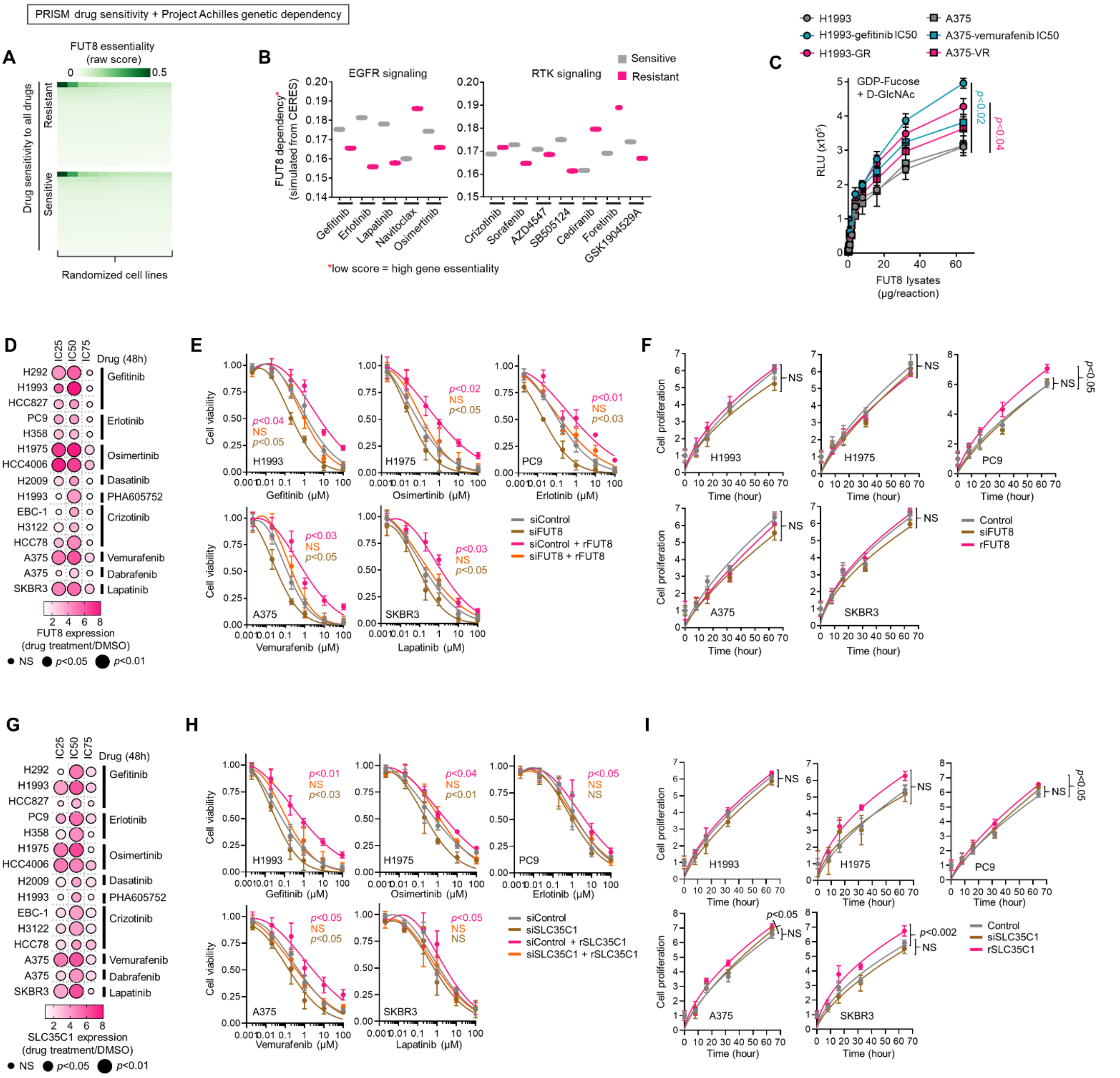


**Supplemental Fig. 8.** FUT8 and SLC35C1 confer resistance to kinase inhibitors in sensitive cancer cells. (**A**) Heatmap visualization of raw essentiality scores of FUT8 in all cell lines screened in both PRISM and Achilles projects. Drug sensitivity of cell lines were binarized and grouped as shown.

(**B**) Mean dependency scores of FUT8 in cell lines indicated as either sensitive or resistant to indicated drugs.

(**C**) GDP-Fuc activity analysis of FUT8 in FUT8 immunoprecipitates from indicated DMSO or drug-treated cell lysates or DR clone lysates. Values indicate luminescence units and are relative to control reaction (means ± SD of three biological replicates). Representative of two independent experiments. For statistical analysis, two-tailed Mann–Whitney *U* test was used.

(**D**) qPCR analysis of FUT8 expression in indicated cells treated with or without IC25, IC50, or IC75 of indicated drugs for 72 h. Values are relative to DMSO and were normalized to GAPDH levels (means ± SD of three biological replicates). *P* values are indicated as size of the corresponding circle; Student's t-test. NS, not significant.

(**E**) Cell viability of indicated cells upon FUT8 RNAi subsequently transfected with or without FUT8 cDNA for 48 h. Cells were treated with indicated drug concentrations for 48 h. Values are relative to DMSO (means ± SD of three biological replicates). Representative of two independent experiments. For statistical analysis, Student's *t*-test was used. NS, not significant.

(**F**) Proliferation of indicated cells upon FUT8 RNAi or FUT8 cDNA transfection as in E. Values are relative to time point 0 (means ± SD of three biological replicates). Representative of two independent experiments. For statistical analysis, Student's *t*-test was used. NS, not significant.

(**G**) qPCR analysis of SLC35C1 expression in indicated cells treated with or without IC25, IC50, or IC75 of indicated drugs for 72 h. Values are relative to DMSO and were normalized to GAPDH levels (means ± SD of three biological replicates). *P* values are indicated as size of the corresponding circle; Student's t-test. NS, not significant.
(**H**) Cell viability of indicated cells upon SLC35C1 RNAi subsequently transfected with or without SLC35C1 cDNA for 48 h. Cells were treated with indicated drug concentrations for 48 h. Values are relative to DMSO (means ± SD of three biological replicates). Representative of two independent experiments. For statistical analysis, Student's *t*-test was used. NS, not significant.
(**I**) Proliferation of indicated cells upon SLC35C1 RNAi or SLC35C1 cDNA transfection as in H. Values are relative to time point 0 (means ± SD of three biological replicates). Representative of two independent experiments. For statistical analysis, Student's *t*-test was used. NS, not significant.

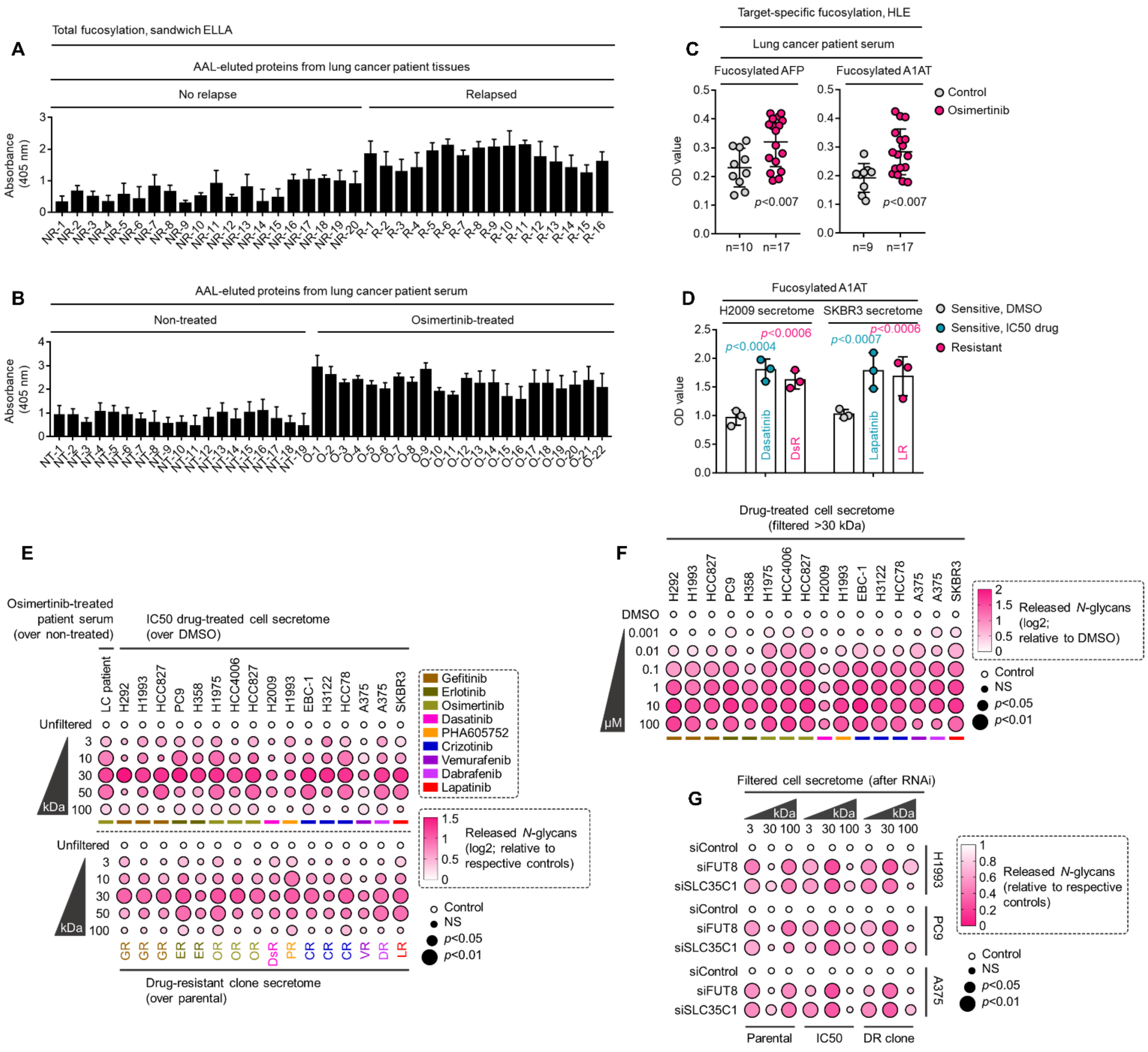


**Supplemental Fig. 9.** Therapy resistance-associated core fucosylation of proteins between 30 and 100 kDa is regulated by FUT8 or SLC35C1.

(**A**) Characterization of fucosylation by sandwich ELLA in indicated tumor tissues from patients with lung cancer who underwent first-line therapy. Values indicate mean absorbance at 405 nm (means ± SD of three biological replicates). Representative of two independent experiments.

(**B**) Characterization of fucosylation by sandwich ELLA in indicated crude sera from patients with lung cancer treated with or without osimertinib. Values indicate mean absorbance at 405 nm (means ± SD of two biological replicates). Representative of two independent experiments.

(**C**) HLE analysis of AFP or A1AT fucosylation in sera from patients with lung cancer treated with or without osimertinib. Values indicate mean absorbance at 450 nm from three replicates (for each sample). Representative of two independent experiments. For statistical analysis, two-tailed Mann–Whitney *U* test was used.

(**D**) HLE analysis of A1AT fucosylation in indicated secretomes derived from sensitive cells treated with or without lapatinib or dasatinib IC50 or indicated DR clones. Values represent mean absorbance at 450 nm (means ± SD of three biological replicates). Representative of two independent experiments. For statistical analysis, two-tailed Mann–Whitney *U* test was used.

(**E**) N-glycan release assay using PNGase F in indicated secretomes from sensitive cells or DR clones following treatment with or without respective drug IC50s for 48 h; or sera from patients treated with or without osimertinib. Cell secretomes were prepared as in Fig. 1H while patient sera were prepared as in Fig. 1F; except filtered according to their indicated nominal molecular weight limit (NMWL). Values indicate mean absorbance at 584 nm and are relative to unfiltered secretome/sera (means ± SD of three biological replicates). Representative of two independent experiments. *P* values are indicated as size of the corresponding circle; Student's *t*-test. NS, not significant.

(**F**) N-glycan release assay using PNGase F in indicated secretomes from sensitive cells following treatment with or without indicated drugs for 48 h. Cell secretomes were prepared as in Fig. 1H; except filtered with a >30 kDa nominal molecular weight limit (NMWL). Values indicate mean absorbance at 584 nm and are relative to DMSO (means ± SD of three biological replicates). Representative of two independent experiments. *P* values are indicated as size of the corresponding circle; Student's *t*-test. NS, not significant.

(**G**) N-glycan release assay using PNGase F in indicated secretomes from sensitive cells or DR clones upon FUT8 or SLC35C1 RNAi for 48 h following treatment with or without drug IC50 (gefitinib for H1993, erlotinib for PC9, and vemurafenib for A375) for 48 h. Cell secretomes were prepared as in Fig. 1H; except filtered according to their indicated nominal molecular weight limit (NMWL). Values indicate mean absorbance at 584 nm and are relative to siControl (means ± SD of three biological replicates). Representative of two independent experiments. *P* values are indicated as size of the corresponding circle; Student's *t*-test. NS, not significant.

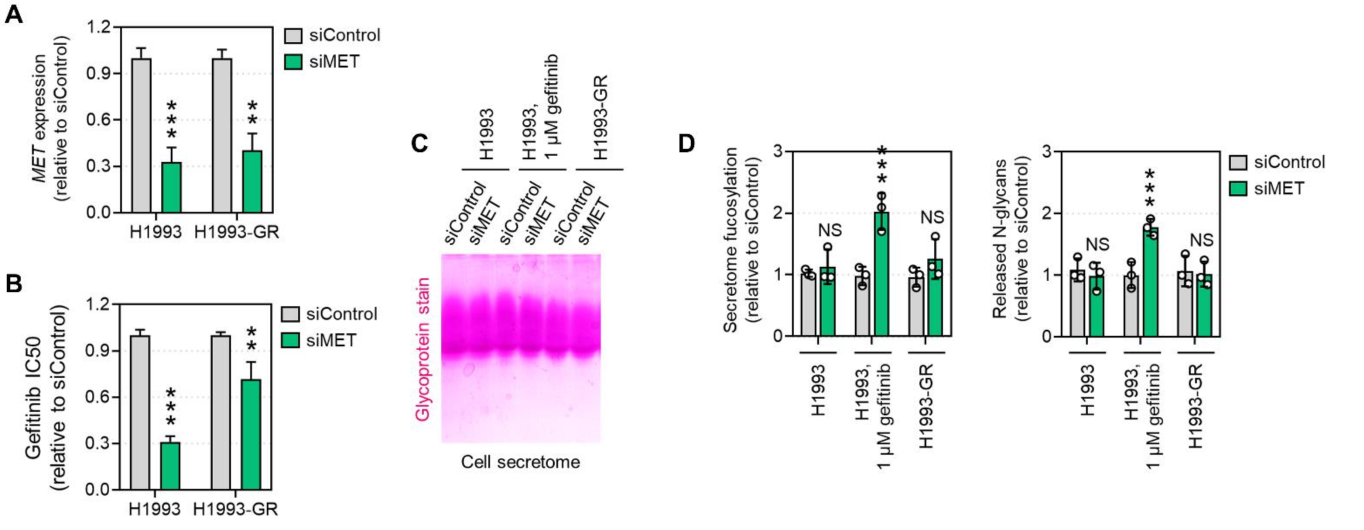


**Supplemental Fig. 10.** MET knockdown in high MET-expressing H1993 cells, but not in GR clones, augments EGFR-TKI-induced secretome fucosylation.

(**A**) qPCR analysis of MET gene expression in indicated cells upon MET RNAi for 48 h. Values are relative to siControl and were normalized to GAPDH levels (means ± SD of three biological replicates). ****P*<0.001, Student's *t*-test.

(**B**) Gefitinib sensitivity assay in indicated cells upon MET RNAi for 48 h. Cells were treated with or without drugs for 72 hours with a concentration dilution series and were assayed for SRB. Values are relative to siControl (means ± SD of two biological replicates). ***P*<0.01, ****P*<0.001, Student's *t*-test.

(**C**) Representative glycoprotein staining of indicated secretomes from H1993 cells or GR clones upon MET RNAi for 48 h followed by treatment with or without 1 μM gefitinib for 48 h.

(**D**) Characterization of fucosylation by sandwich ELLA and N-glycan release assay in indicated cells as in C. Values are relative to siControl (means ± SD of three biological replicates). **P*<0.05, ***P*<0.01, ****P*<0.001, Student's *t*-test. NS, not significant.

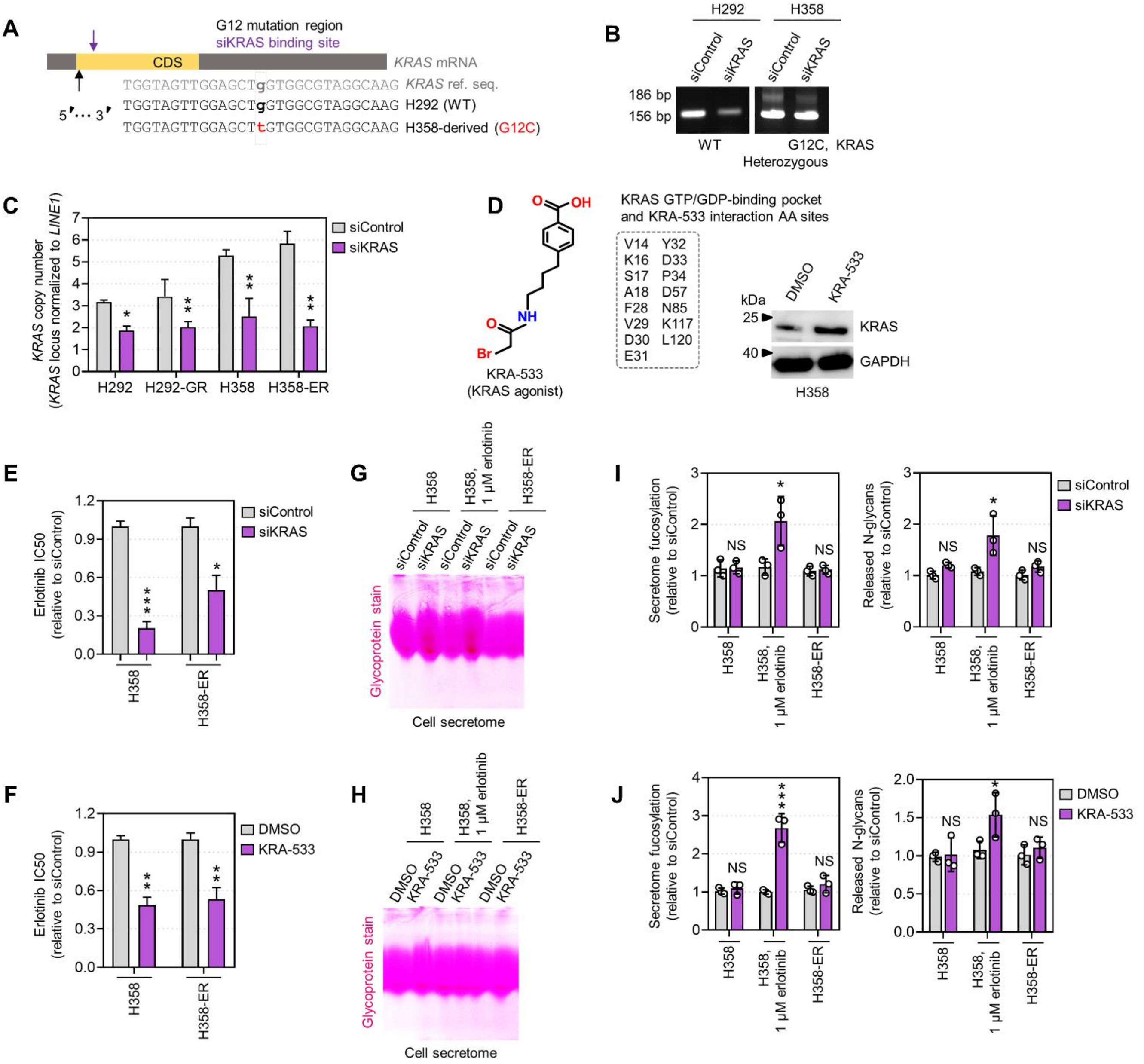


**Supplemental Fig. 11.** Inhibition of oncogenic KRAS in mutant H358 cells, but not in ER clones, promotes EGFR-TKI-induced secretome fucosylation.

(**A**) Binding sites of KRAS siRNA sequence (violet) used in this study in close proximity to the region of G12 (and G13) codon mutations demonstrated in the coding sequence (CDS) of a mature KRAS mRNA map. H292, harboring wild type (WT) KRAS, was used as reference.

(**B**) Visualization of KRAS WT and mutant transcripts of H292 and H358 cells, respectively, following KRAS amplification with primers designed to introduce base substitution creating a BstNI recognition site specific for the WT codon 12 but not for the codon with KRAS mutation. BstNI digestion cuts the WT allele (in H292 cells) to produce a 156 bp fragment whereas the mutant allele remains uncut to produce a 186 bp fragment (in H358 cells). Note that the both WT and mutant bands were observed in H358 cells as they harbor a heterozygous G12 codon mutation.

(**C**) KRAS copy number analysis in KRAS mutant H358 cells and WT H292 cells and their respective DR clones upon KRAS RNAi for 60 h. Values are relative to siControl and were normalized to LINE1 elements (means ± SD of three biological replicates). *$P$<0.05, **$P$<0.01, Student's $t$-test.

(**D**) Structure of the small molecule NSC112533 (KRA-533) and its known amino acid interaction sites with the GTP/GDP-binding pocket of the KRAS protein. Beside shows a Western blot analysis of KRAS expression in H358 cells after treatment with or without 8 μM KRA-533 for 48 h. GAPDH was used as a loading control. Representative of two independent experiments.

(**E**) Erlotinib sensitivity assay in indicated cells upon KRAS RNAi for 48 h. Cells were treated with or without drugs for 72 hours with a concentration dilution series and were assayed for SRB. Values are relative to

siControl (means ± SD of three biological replicates). *$P<0.05$, ***$P<0.001$, Student's *t*-test.
(**F**) Erlotinib sensitivity assay in indicated cells after treatment with or without 8 μM KRA-533 for 48 h. Cells were treated with or without drugs for 72 hours with a concentration dilution series and were assayed for SRB. Values are relative to DMSO (means ± SD of three biological replicates). **$P<0.01$, Student's *t*-test.
(**G**) Representative glycoprotein staining of indicated secretomes from H358 cells or ER clones upon KRAS RNAi for 48 h followed by treatment with or without 1 μM erlotinib for 48 h.
(**H**) Representative glycoprotein staining of indicated secretomes from H358 cells or ER clones after treatment with or without 8 μM KRA-533 for 48 h and with or without 1 μM erlotinib for 48 h.
(**I**) Characterization of fucosylation by sandwich ELLA and N-glycan release assay in indicated cells as in G. Values are relative to siControl (means ± SD of two biological replicates). *$P<0.05$, Student's *t*-test. NS, not significant.
(**J**) Characterization of fucosylation by sandwich ELLA and N-glycan release assay in indicated cells as in H. Values are relative to DMSO (means ± SD of two biological replicates). *$P<0.05$, ***$P<0.001$, Student's *t*-test. NS, not significant.

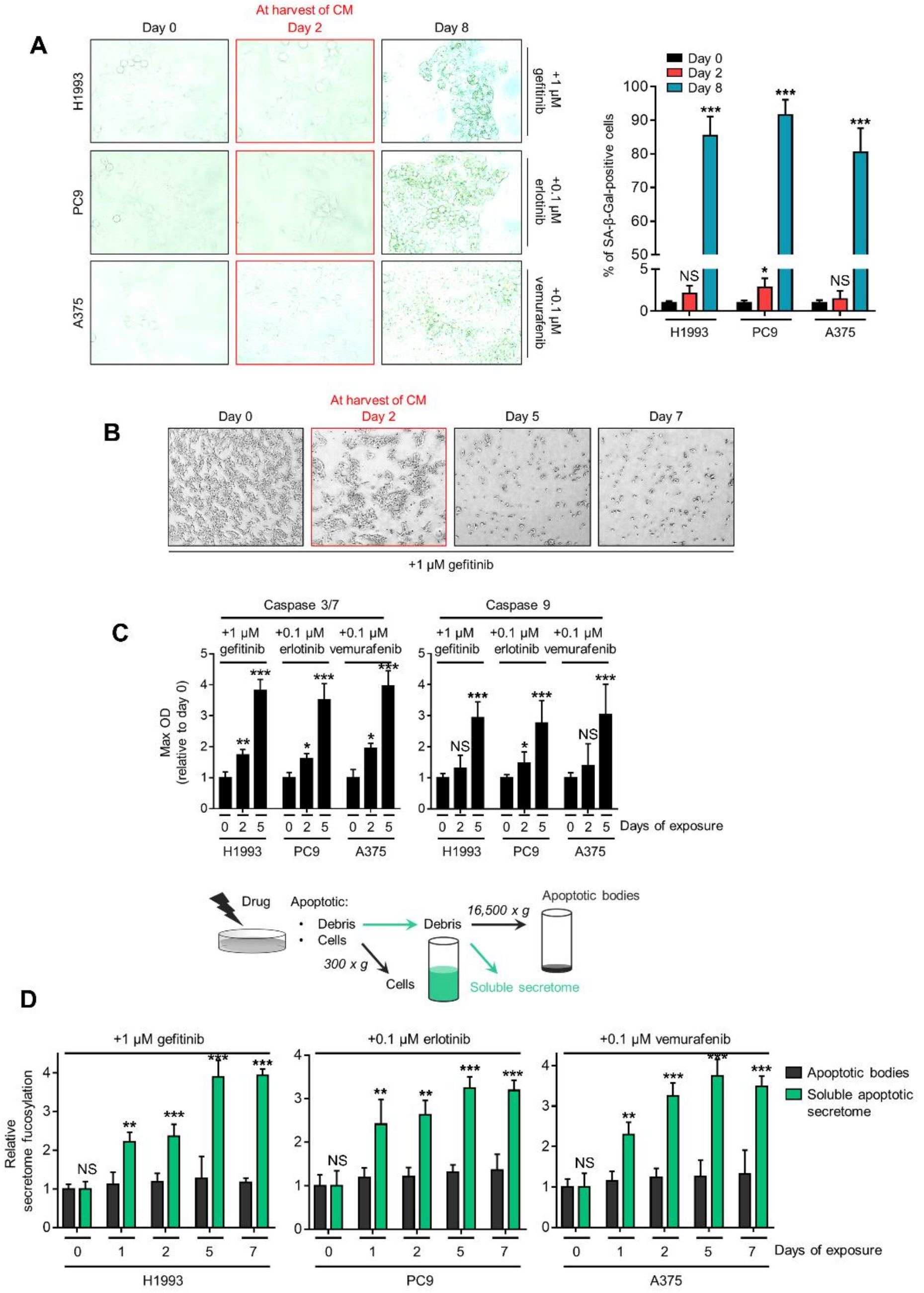


**Supplemental Fig. 12.** Drug-induced secretome fucosylation occurs before apoptosis and senescence.
(**A**) SA-β-gal staining of indicated cells treated with indicated drug concentrations for 2 or 8 days. Cells were passaged over ~8 times before the experiment. Representative images are shown (left) along with quantified data as percentage (right; means ± SD of three biological replicates). **P*<0.05, ****P*<0.001, Student's *t*-test. NS, not significant.
(**B**) Morphological evaluation of H1993 cells treated with 1 μM gefitinib for indicated times.
(**C**) Caspase 3/7 DEVDase and caspase 9 activities of indicated cells and treatment conditions as in A. Values are relative to time point 0 (means ± SD of two biological replicates). **P*<0.05, ***P*<0.01, ****P*<0.001, Student's *t*-test. NS, not significant.
(**D**) Sandwich ELLA of apoptotic cell debris and secretome from indicated cells treated with indicated drug concentrations for indicated times by means of centrifugation. Samples were prepared as in the top panel. Values are relative to time point 0 (means ± SD of two biological replicates). ***P*<0.01, ****P*<0.001, Student's *t*-test. NS, not significant.

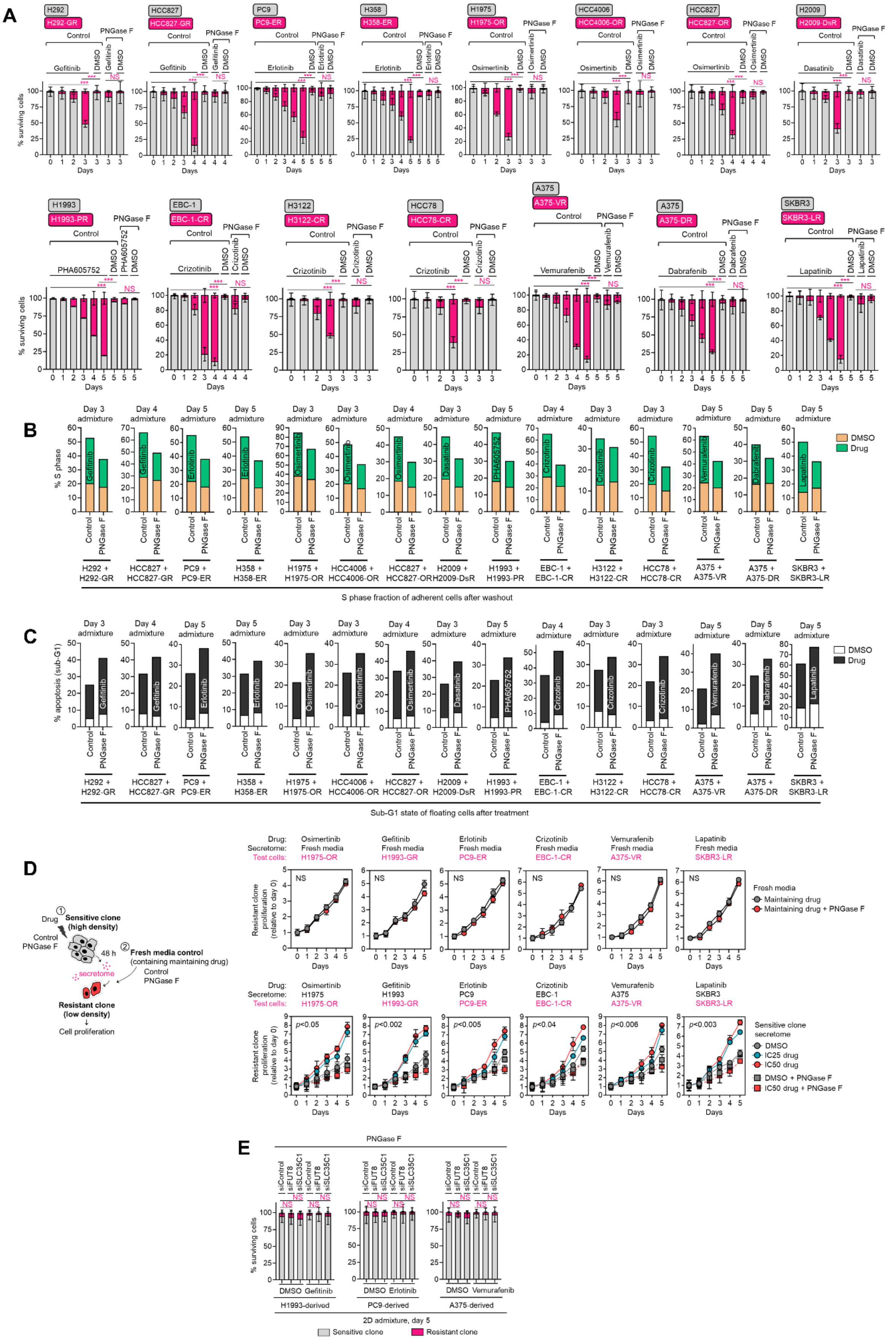

**Supplemental Fig. 13.** Resistance rebound and apoptosis in 2D admixtures.
(**A**) Tracking of both fluorescently-tagged cells in 2D cell admixtures prepared as in Fig. 2A, treated with or without indicated drugs [gefitinib (1 μM in H292, 0.08 μM in HCC827), erlotinib (0.1 μM in PC9, 1 μM in H358), osimertinib (0.08 μM in H1975, 0.01 μM in HCC4006, 0.08 μM in HCC827), dasatinib (0.8 μM in H2009), PHA665752 (0.6 μM in H1993), crizotinib (0.08 μM in EBC-1, 0.5 μM in H3122, 1 μM in HCC78), vemurafenib (0.1 μM in A375), dabrafenib (0.03 μM in A375), lapatinib (0.4 μM in SKBR3)], and incubated with or without

10 μg/mL recombinant PNGase F for indicated times. Values are relative to day 0 (means ± SD of three biological replicates). *$P$<0.05, **$P$<0.01, ***$P$<0.001, two-tailed Mann–Whitney $U$ test. NS, not significant.
(**B** and **C**) Cell cycle states of adherent cells and apoptosis of floating cells in indicated cell admixtures with same conditions as in A at indicated times. Representative of two independent experiments.
(**D**) Proliferation of indicated DR clones prepared as in the CM co-culture schematic. Cell admixtures were incubated with or without 10 μg/mL recombinant PNGase F for indicated times. Maintaining drug concentrations are shown in Supplemental Fig. 6A. Values are relative to time point 0 (means ± SD of three biological replicates). For statistical analysis, Student's $t$-test was used. NS, not significant.
(**E**) Similar tracking experiments as in A, except upon FUT8 or SLC35C1 RNAi in sensitive cells for 48 h prior to admixing and culture for 5 days. Cell admixtures treated with or without indicated drugs as in A and were incubated with or without 10 μg/mL recombinant PNGase F for indicated times. Values are relative to day 0 (means ± SD of two biological replicates). For statistical analysis, two-tailed Mann–Whitney $U$ test was used. NS, not significant.

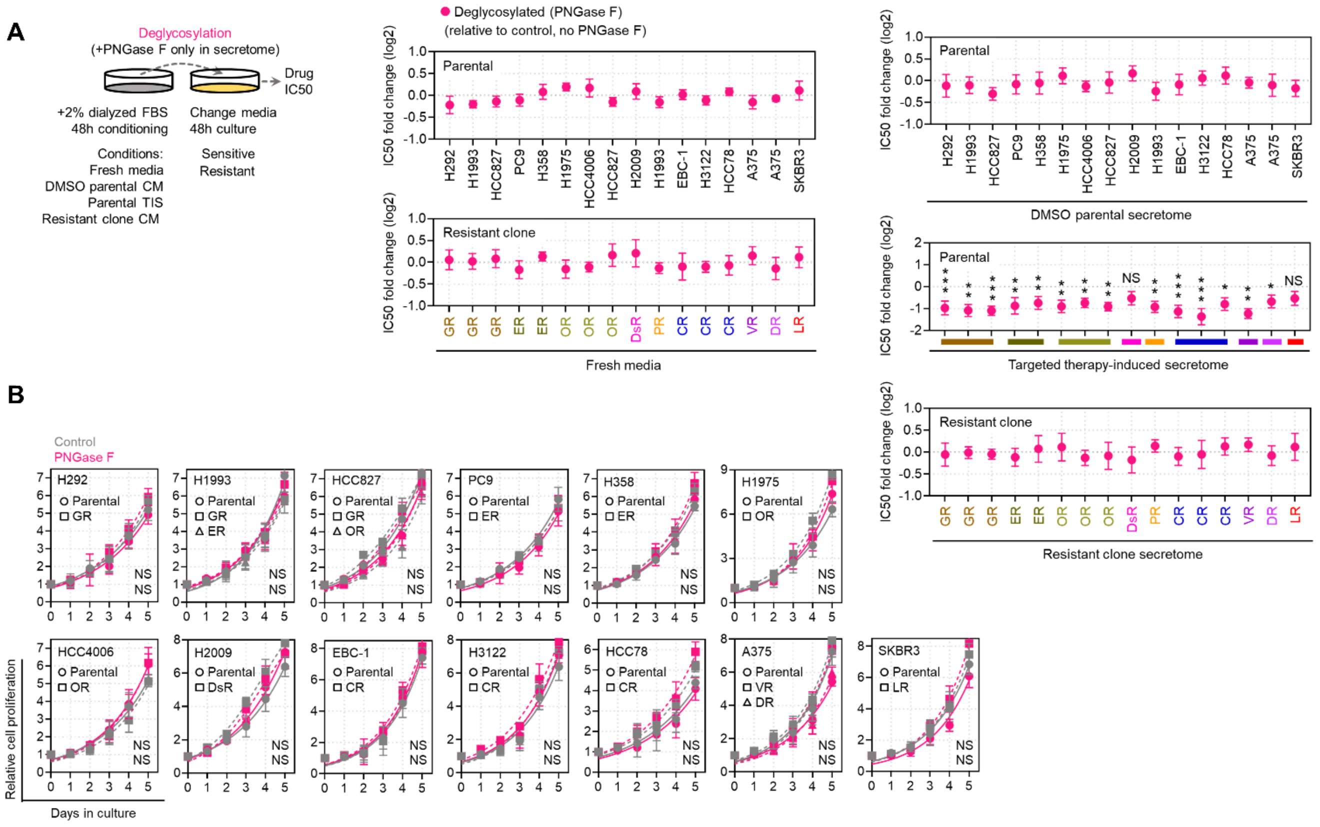

**Supplemental Fig. 14.** Effects of de-N-glycosylation in cell cultures.

(**A**) Drug sensitivity assays in indicated sensitive cells or DR clones prepared as in the schematic. Prior to CM co-culture, secretomes were exogenously treated with or without 8U PNGase F. Cells were treated with or without drugs for 72 hours with a concentration dilution series and were assayed for SRB. Values are relative to DMSO (means ± SD of three biological replicates). **P*<0.05, ***P*<0.01, ****P*<0.001, Student's *t*-test. NS, not significant.

(**B**) Proliferation of indicated sensitive or DR clones treated with or without 10 μg/mL recombinant PNGase F for indicated times. Values are relative to time point 0 (means ± SD of three biological replicates). For statistical analysis, Student's t-test was used. NS, not significant.

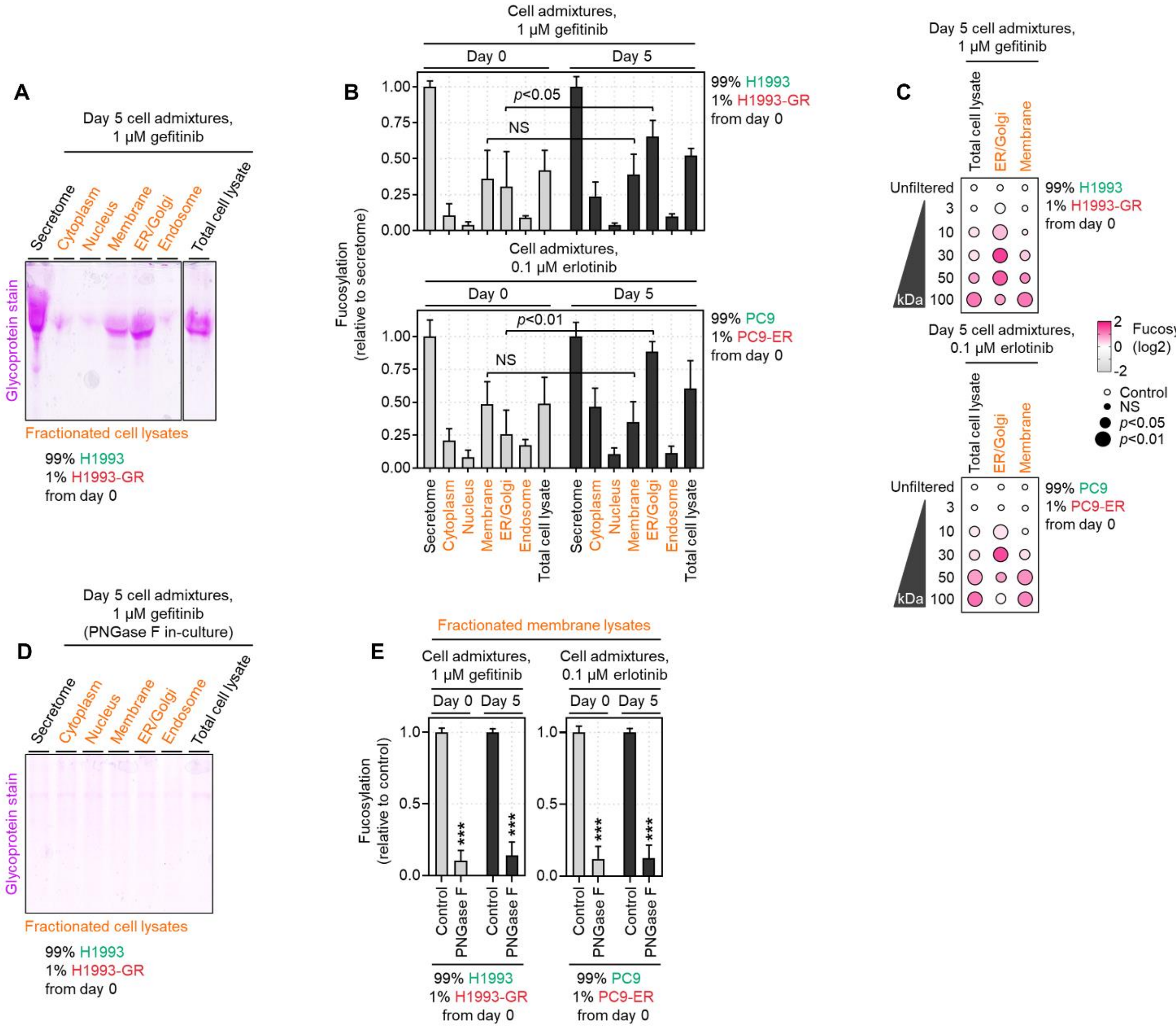


**Supplemental Fig. 15.** Cell membrane-bound N-glycoproteins are dispensable in promoting drug-resistant population expansion in drug-treated admixture cultures.

(**A**) Representative glycoprotein staining of H1993 cell admixtures at day 5 prepared as in Fig. 2A, treated with 1 μM gefitinib. Cell secretomes, total cell lysates, and pooled subcellular fraction lysates were collected from cell admixtures.

(**B**) Sandwich ELLA of indicated H1993 and PC9 cell admixtures at days 0 and 5 prepared as in Fig. 2A, treated with 1 μM gefitinib and 0.1 μM erlotinib, respectively. Cell secretomes, total cell lysates, and pooled subcellular fraction lysates were collected from cell admixtures. Values are relative to cell secretome (means ± SD of three biological replicates). For statistical analysis, Student's *t*-test was used. NS, not significant.

(**C**) Sandwich ELLA of indicated H1993 and PC9 cell admixtures as in B, except filtered according to their indicated nominal molecular weight limit (NMWL). Values are relative to unfiltered lysates (means ± SD of three biological replicates). *P* values are indicated as size of the corresponding circle; Student's *t*-test. NS, not significant.

(D) Representative glycoprotein staining of H1993 cell admixtures as in A, except incubated with 10 μg/mL recombinant PNGase F. Cell secretomes, total cell lysates, and pooled subcellular fraction lysates were collected from cell admixtures.

(E) Sandwich ELLA of indicated H1993 and PC9 cell admixtures as in B, except incubated with 10 μg/mL recombinant PNGase F. Only results from pooled subcellular membrane fraction lysates collected from cell admixtures are shown. ***$P<0.001$, Student's *t*-test.

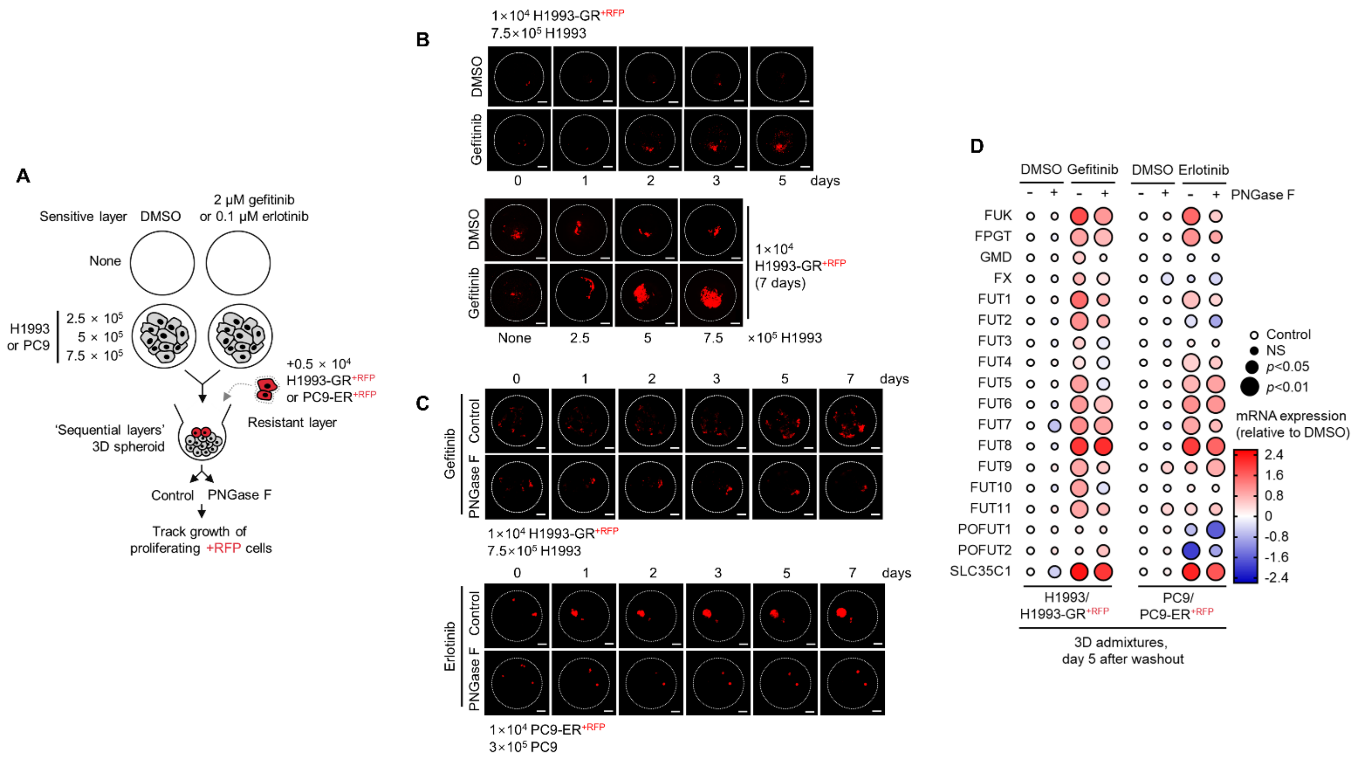


**Supplemental Fig. 16.** Outgrowth of DR clones in regressing 3D spheroids is associated with expression of fucose salvage genes.

(**A**) Schematic of 'sequential layer' admixture culture.

(**B**) Representative confocal images of indicated RFP-labeled GR clones in 3D spheroid admixtures prepared as in A and treated with or without 2 μM gefitinib for indicated times. Scale bar indicates 100 μm. Representative of two independent experiments.

(**C**) Representative confocal images of indicated RFP-labeled GR clones in 3D spheroid admixtures prepared as in A, treated with either 2 μM gefitinib or 0.1 μM erlotinib, and incubated with or without 10 μg/mL recombinant PNGase F for indicated times. Scale bar indicates 100 μm. Representative of two independent experiments.

(**D**) qPCR analysis of indicated gene expression in 3D cell admixtures prepared as in A, treated with or without 2 μM gefitinib or 0.1 μM erlotinib, and incubated with or without 10 μg/mL recombinant PNGase F for 5 days. Values are relative to DMSO and were normalized to GAPDH levels (means ± SD of three biological replicates). *P* values are indicated as size of the corresponding circle; Student's *t*-test. NS, not significant.

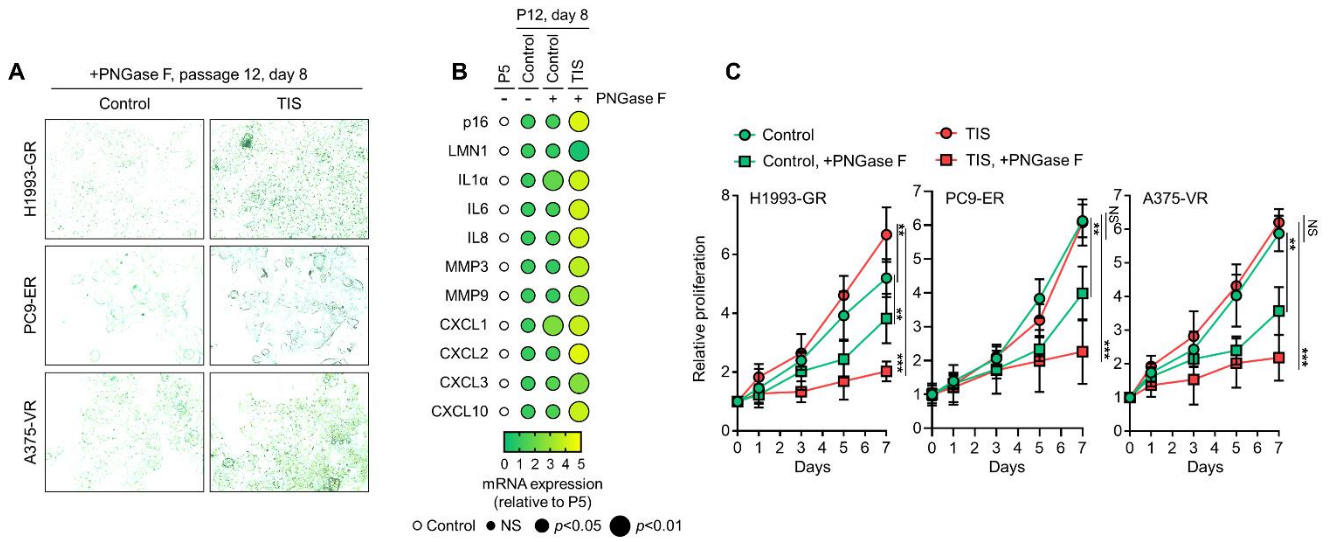


**Supplemental Fig. 17.** TIS de-N-glycosylation promotes senescence in long-term grown DR clones.

(**A**) SA-β-gal staining of indicated DR clones incubated with or without indicated TIS (secretomes derived from parental cells treated 1 μM gefitinib, 0.1 μM erlotinib, or 0.1 μM vemurafenib for 2 days) and with 10 μg/mL recombinant PNGase F for 8 days. Representative of two independent experiments.

(**B**) qPCR analysis of indicated SASP gene expression in GR clones as in A. Values are relative to P5 and were normalized to GAPDH levels (means ± SD of three biological replicates). *P* values are indicated as size of the corresponding circle; Student's *t*-test. NS, not significant.

(**C**) Proliferation of indicated DR clones as in A. Values are relative to time point 0 (means ± SD of three biological replicates). *$P<0.05$, **$P<0.01$, ***$P<0.001$, Student's *t*-test. NS, not significant.

**A**

Label-free fucosylated secretome analysis (30-70 kDa fucosylated proteins)

| Group | | MS/MS identified | | | Isotope patterns sequenced ($z$>1) | Peptide sequence identified |
|---|---|---|---|---|---|---|
| | | SIL | PEAK | % | | |
| H1993 | Technical replicate #1 | 1121 | 139 | 1.7 | 26825 | 411 |
| | Technical replicate #2 | 1092 | 125 | 1.6 | 26687 | 474 |
| H1993, 1 μM gefitinib | Technical replicate #1 | 1135 | 96 | 1.8 | 26258 | 450 |
| | Technical replicate #2 | 1144 | 127 | 1.6 | 28935 | 462 |
| H1993-GR | Technical replicate #1 | 1134 | 112 | 1.7 | 29145 | 503 |
| | Technical replicate #2 | 1139 | 116 | 1.6 | 28939 | 518 |

**B**

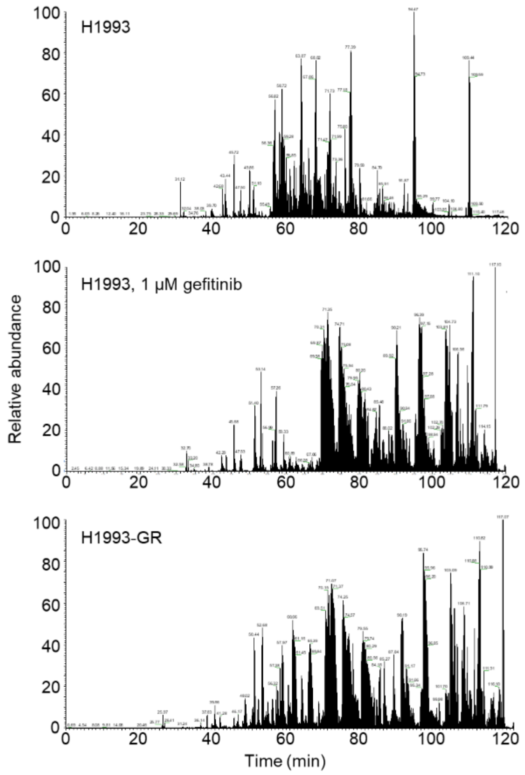


**C**

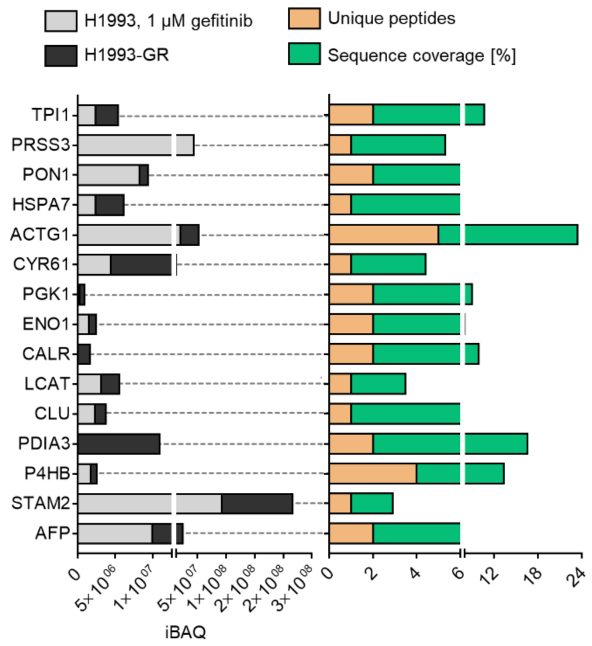


**Supplemental Fig. 18.** In-gel N-glycome analysis.

(**A**) Summary information on label-free secretome analysis.

(**B**) Representative base peak chromatogram of protein digest samples from indicated cells or DR clone by a 120 min LC gradient. Selected chromatographic peaks are labeled with m/z of the underlying peptide. Representative profile of two replicates.

(**C**) iBAQ intensities (relative protein abundances), unique peptides, and sequence coverage in percentile of indicated proteins in secretomes of gefitinib-treated H1993 cells or GR clone. Top 16 protein hits with MWs between 30 and 70 kDa are shown. Results were analyzed from two biological replicates.

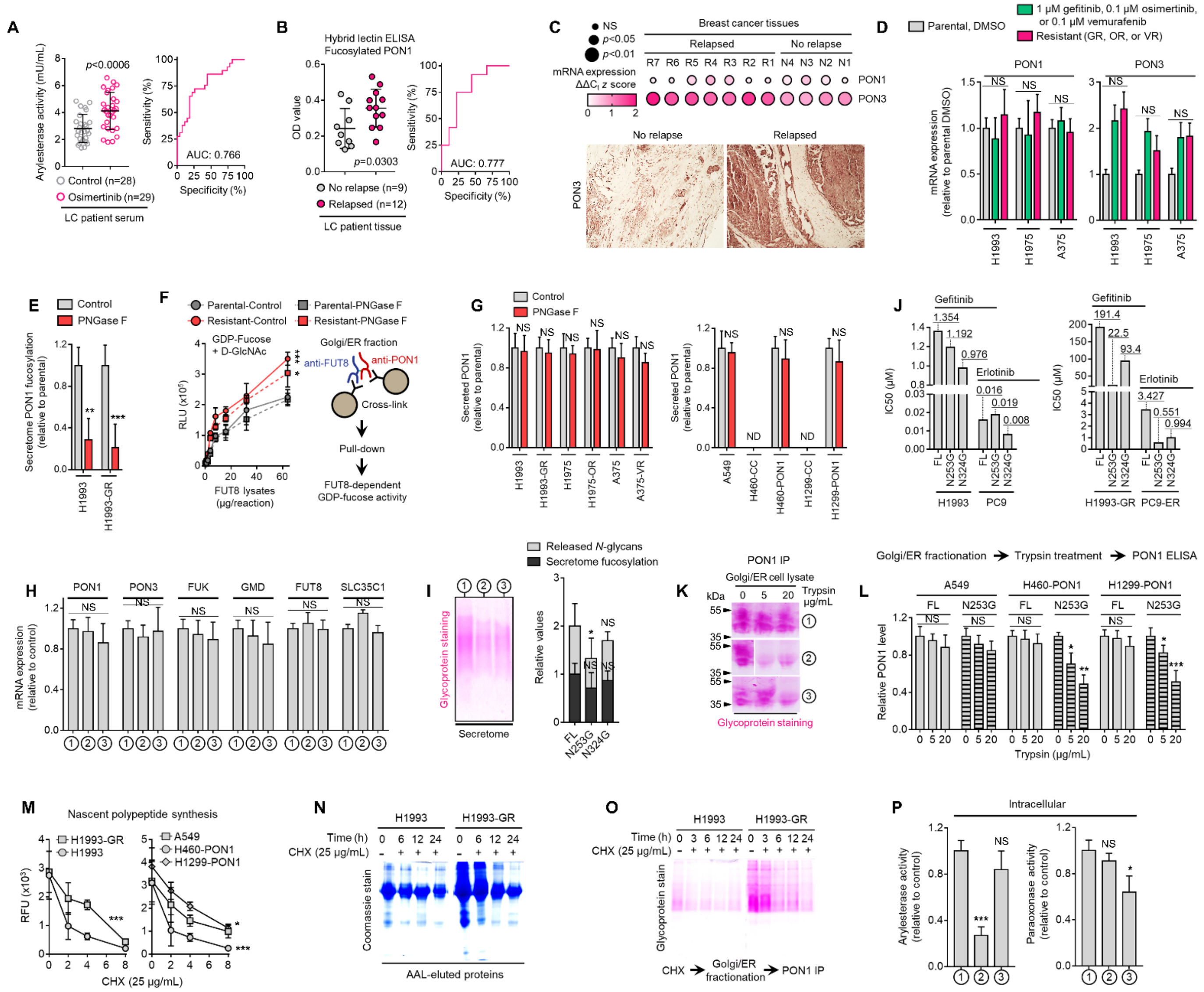

**Supplemental Fig. 19.** Core fucosylation directs PON1 secretory process in DR clones and PON1-overexpressing cells.

(**A**) Quantification of arylesterase activity in indicated crude patient sera. Values indicate mean absorbance at 217 nm from three replicates. Representative of two independent experiments. ROC curve for PON1 arylesterase activity is also shown. For statistical analysis, two-tailed Mann–Whitney *U* test was used.

(**B**) HLE analysis of PON1 fucosylation in indicated tumor tissues from patients with lung cancer who underwent first-line therapy. Representative of two independent experiments. ROC curve for PON1 fucosylation is also shown. For statistical analysis, two-tailed Mann–Whitney *U* test was used.

(**C**) qPCR analysis of PON1 and PON3 expression in indicated FFPE tumor tissue sections from patients with breast cancer who underwent sequential multidrug therapy. Log-transformed gene expression values are relative to the non-relapsed sample with lowest expression for the indicated gene (not displayed in the heatmap) and were normalized to GAPDH levels (means ± SD of three biological replicates). *P* values are indicated as size of the corresponding circle; Student's *t*-test. NS, not significant. Bottom panel shows PON3 immunohistochemical analysis of indicated tumor sections. All sections were photographed with an inverted phase-contrast microscope (original magnification, ×200). Representative of two independent experiments.

(**D**) qPCR analysis of PON1 and PON3 expression in indicated cells or DR clones. Values are relative to DMSO or parental (means ± SD of two biological replicates). For statistical analysis, Student's t-test was used. NS, not significant.

(**E**) HLE analysis of PON1 fucosylation in secretomes derived from cells or GR clone incubated with or without 10 μg/mL recombinant PNGase F for 48 h. Values are relative to non-treated (means ± SD of two biological replicates). $^{**}P<0.01$, $^{***}P<0.001$, Student's *t*-test. NS, not significant.

(**F**) GDP-Fuc activity analysis of FUT8 in cross-linked FUT8 and PON1 co-immunoprecipitates from Golgi/ER fractionated H1993 or H1993-GR lysates. Prior to fractionation, cells were incubated with or without 10 μg/mL recombinant PNGase F for 48 h. Values indicate luminescence units and are relative to control reaction

(means ± SD of three biological replicates). **P*<0.05, ****P*<0.001, two-tailed Mann–Whitney *U* test.
(**G**) ELISA analysis of PON1 in secretomes derived from sensitive cells, DR clones, WT or PON1-edited cells. Cells were incubated with or without 10 μg/mL recombinant PNGase F for 48 h. Values are relative to non-treated (means ± SD of three biological replicates). For statistical analysis, Student's *t*-test was used. NS, not significant.
(**H**) qPCR analysis of indicated gene expression in indicated sensitive cells or DR clones. Cells were transfected with PON1-FL or mutant PON1 constructs for 36 h. Values are relative to PON1-FL (means ± SD of three biological replicates). For statistical analysis, Student's *t*-test was used. NS, not significant.
(**I**) Characterization of secretome fucosylation by glycoprotein staining, sandwich ELLA, and N-glycan release assay in H1993-GR transfected with PON1-FL or mutant PON1 constructs for 36 h. Values are relative to PON1-FL (means ± SD of three biological replicates). **P*<0.05, Student's *t*-test. NS, not significant.
(**J**) Drug sensitivity assays in indicated sensitive cells or DR clones. Cells were transfected with PON1-FL or mutant PON1 constructs for 36 h and treated with or without indicated drugs for 72 hours with a concentration dilution series and were assayed for SRB. Representative of three independent experiments.
(**K**) Glycoprotein staining of PON1 immunoprecipitates from Golgi/ER fractionated H1993-GR lysates upon exogenous treatment with or without indicated trypsin concentrations. Prior to fractionation, cells were transfected with PON1-FL or mutant PON1 constructs for 36 h. Representative of two independent experiments.
(**L**) ELISA analysis of PON1 expression in indicated WT or PON1-edited cells upon transfections with PON1-FL or mutant PON1 constructs for 36 h. Golgi/ER fractionated cell lysates were exogenously treated with or without indicated trypsin concentrations. Values are relative to no treatment (means ± SD of three biological replicates). **P*<0.05, ***P*<0.01, ****P*<0.001, Student's *t*-test. NS, not significant.
(**M**) EZClick labeling analysis of polypeptide synthesis in indicated cells, DR clones, WT or PON1-edited cells treated with or without 25 μg/mL CHX for indicated times. Values indicate raw fluorescence units (means ± SD of two biological replicates). **P*<0.05, ****P*<0.001, Student's *t*-test. NS, not significant.
(**N**) Representative Coomassie stained SDS-PAGE gels showing fucosylated secretome proteins from indicated cells or GR clone treated with or without 25 μg/mL CHX for indicated times. Secretomes were concentrated using a >3 kDa NMWL filter.
(**O**) Glycoprotein staining of PON1 immunoprecipitates from Golgi/ER fractionated H1993 or H1993-GR lysates. Prior to fractionation, cells were treated with or without 25 μg/mL CHX for indicated times. Secretomes were concentrated using a >10 kDa NMWL filter. Representative of two independent experiments.
(**P**) Quantification of intracellular arylesterase and paraoxonase activities in H1993-GR transfected with PON1-FL or mutant PON1 constructs. Values indicate mean fluorescence units at 217 and 412 nm, respectively (means ± SD of three biological replicates). **P*<0.05, ****P*<0.001, Student's *t*-test. NS, not significant.

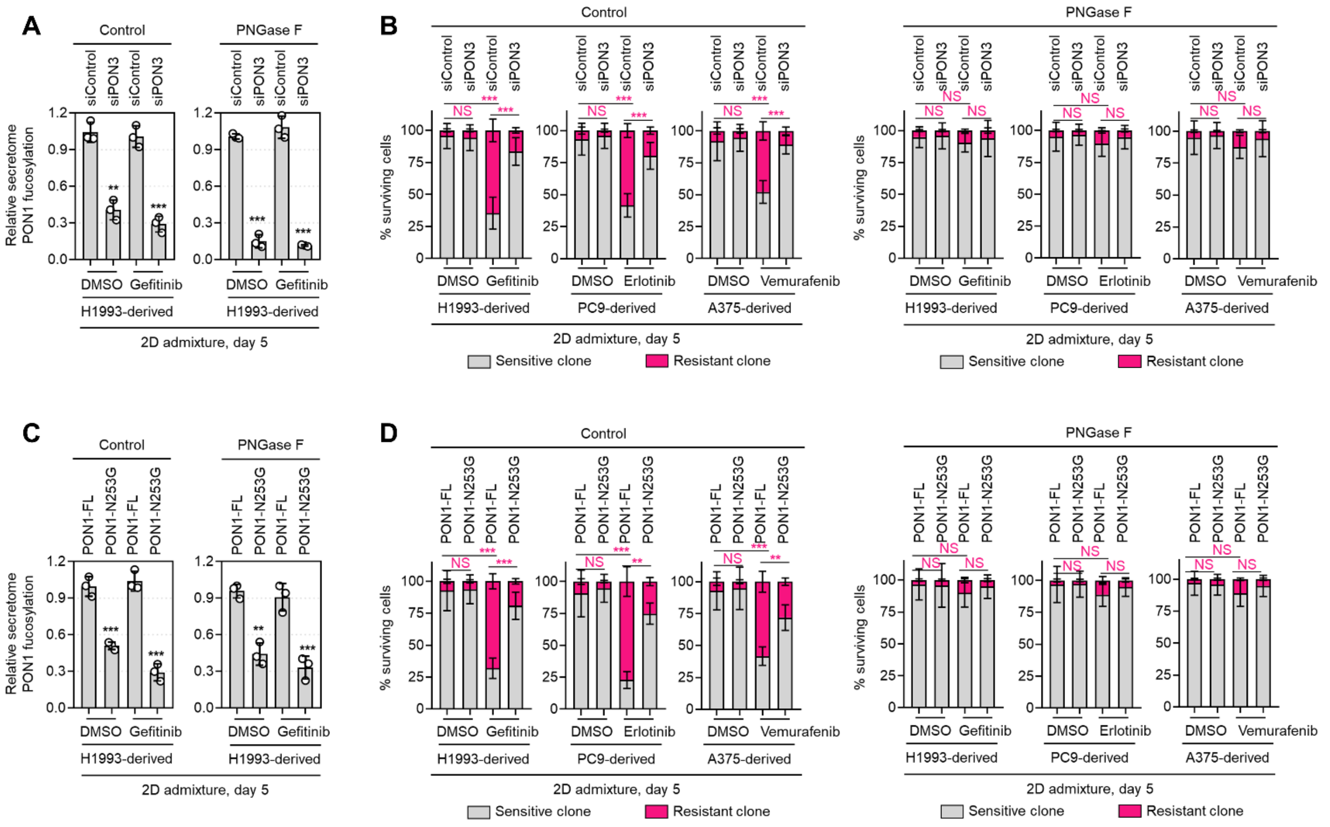


**Supplemental Fig. 20.** PON1 secretome fucosylation promotes TIS-induced resistance rebound in regressing cell admixtures.
(**A**) HLE analysis of secretome PON1 fucosylation in indicated day 5 H1993 cell admixtures prepared as in Fig. 2A. Prior to admixture, sensitive cells were transfected with PON3 RNAi for 48 h. Cell admixtures were treated with or without 1 μM gefitinib and incubated with or without 10 μg/mL recombinant PNGase F. Values are relative to siControl (means ± SD of three biological replicates). ***P*<0.01, ****P*<0.001, Student's *t*-test.
(**B**) Similar tracking experiments as in Fig. 2G, except upon PON3 RNAi in sensitive cells for 48 h prior to admixing and culture for 5 days. H1993 admixture was treated with or without 1 μM gefitinib, PC9 admixture was treated with or without 0.1 μM erlotinib, and A375 admixture was treated with or without 0.1 μM vemurafenib. Cell admixtures were incubated with or without 10 μg/mL recombinant PNGase F.
Values are relative to day 0 (means ± SD of two biological replicates). ****P*<0.001, two-tailed Mann–Whitney *U* test. NS, not significant.
(**C**) HLE analysis of secretome PON1 fucosylation in indicated day 5 H1993 cell admixtures as in A, except upon transfection with full-length PON1 or PON1-N253G construct for 36 h in sensitive cells prior to admixing and culture for 5 days. Cell admixtures were treated with or without 1 μM gefitinib and incubated with or without 10 μg/mL recombinant PNGase F. Values are relative to siControl (means ± SD of three biological replicates). ***P*<0.01, ****P*<0.001, Student's *t*-test.
(**D**) Similar tracking experiments as in Fig. 2G, except upon transfection with full-length PON1 or PON1-N253G construct for 36 h in sensitive cells prior to admixing and culture for 5 days. Cell admixtures were treated with or without 1 μM gefitinib and incubated with or without 10 μg/mL recombinant PNGase F. Values are relative to siControl (means ± SD of three biological replicates). ***P*<0.01, ****P*<0.001, two-tailed Mann–Whitney *U* test. NS, not significant.

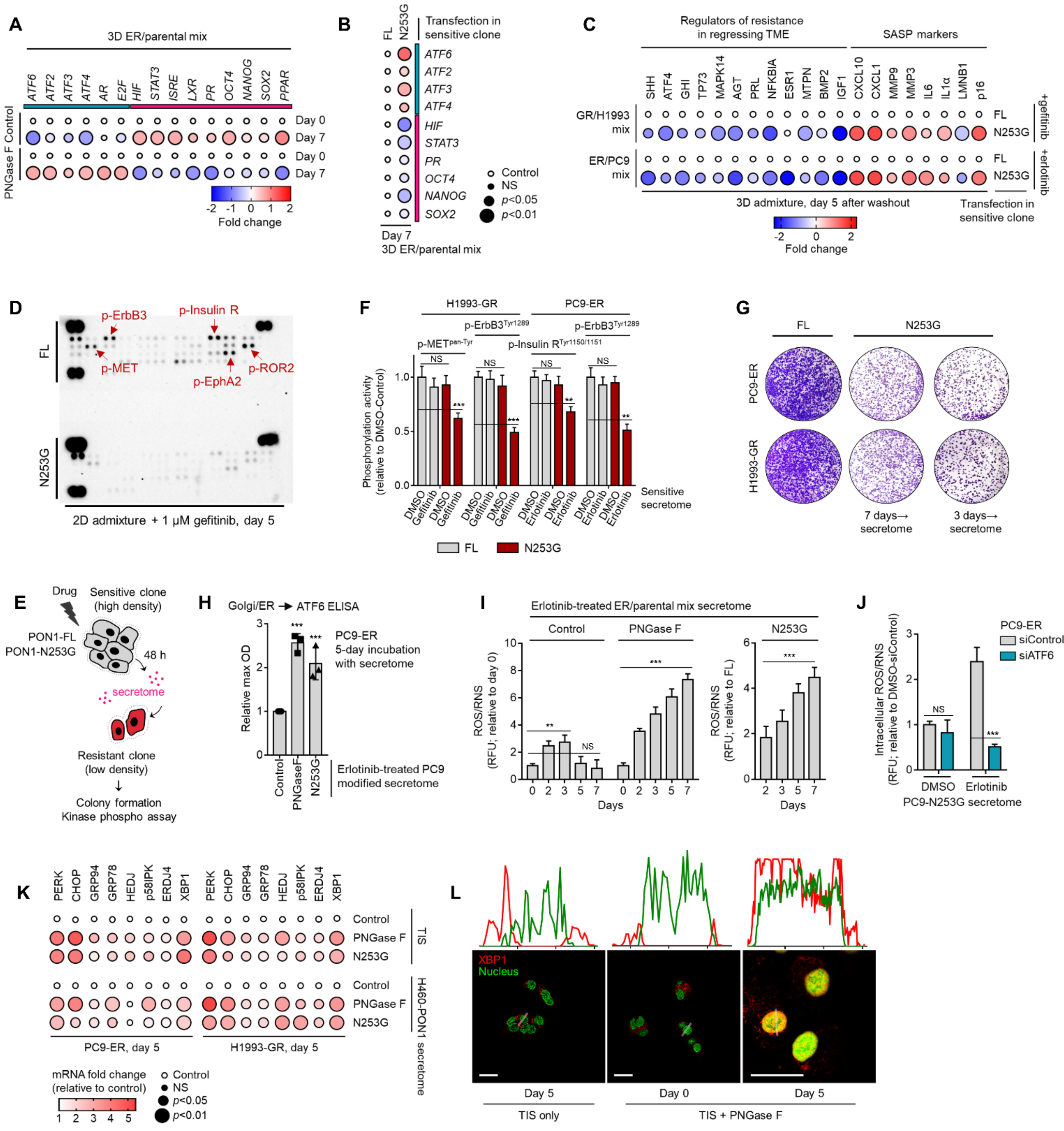


**Supplemental Fig. 21.** Inhibition of TIS-specific PON1 fucosylation prevents resistance rebound via suppression of kinase phospho-proteome and induction of UPR target genes.

(**A**) qPCR analysis of indicated gene expression in 3D cell admixtures with same conditions as in Fig. 5A, except treated with or without 0.1 μM erlotinib for 7 days. Values are relative to day 0 control and were normalized to GAPDH levels (means ± SD of three biological replicates). *P* values are indicated as size of the corresponding circle; Student's *t*-test. NS, not significant.

(**B**) qPCR analysis of indicated gene expression in 3D cell admixtures with same conditions as in A upon transfection with PON1-FL or PON1-N253G construct for 36 h and treatment with or without 0.1 μM erlotinib. Values are relative to FL and were normalized to GAPDH levels (means ± SD of three biological replicates). *P* values are indicated as size of the corresponding circle; Student's *t*-test. NS, not significant.

(**C**) qPCR analysis of indicated gene expression in 3D cell admixtures with same conditions as in Fig. 5A, except transfected with PON1-FL or PON1-N253G construct for 36 h and treated with 2 μM gefitinib or 0.1 μM erlotinib for 5 days. Values are relative to PON1-FL and were normalized to GAPDH levels (means ± SD of three biological replicates). *P* values are indicated as size of the corresponding circle; Student's *t*-test. NS, not significant.

(**D**) Phospho-RTK array of indicated 2D cell admixtures and conditions as in Fig. 2G, except sensitive cells

were transfected with PON1-FL or PON1-N253G construct for 36 h. The blots reflect the phosphorylation status of 49 RTKs. Each RTK is spotted in duplicate, and the three pairs of dots in each corner are positive or negative controls. Representative of two independent experiments.
(**E**) Schematic of CM co-culture.
(**F**) ELISA sandwich-based measurement of indicated RTK phosphorylation in indicated DR clones prepared as in E. Values are relative to DMSO (means ± SD of three biological replicates). **$P$<0.01, ***$P$<0.001, Student's $t$-test. NS, not significant.
(**G**) Colony formation of indicated DR clones prepared as in E. Representative of two independent experiments.
(**H**) ELISA analysis of ATF6 expression in Golgi/ER fractionated PC9-ER grown for 5 days in indicated secretomes from 0.1 µM erlotinib-treated PC9 cells exogenously treated with total 8U PNGase F or transfected with PON1-FL or PON1-N253G construct for 36 h. Values are relative to PON1-FL (means ± SD of three biological replicates). ***$P$<0.001, Student's $t$-test.
(**I**) ROS/RNS detection in secretomes from 0.1 µM erlotinib-treated cell admixtures as in H, except at indicated times. Values are relative to PON1-FL (means ± SD of three biological replicates). **$P$<0.01, ***$P$<0.001, Student's $t$-test. NS, not significant.
(**J**) Intracellular ROS/RNS detection in PC9-ER upon ATF6 RNAi for 48 h and grown in secretomes from PON1-N253G-transfected PC9 cells treated with or without 0.1 µM erlotinib for 72 h. Values are relative to DMSO siControl (means ± SD of two biological replicates). ***$P$<0.001, Student's $t$-test. NS, not significant.
(**K**) qPCR analysis of indicated gene expression in 3D cell admixtures with same conditions as in Fig. 5A, fig. S14A, or admixtures grown in H460-PON1-derived secretomes. Sensitive cells were transfected with PON1-FL or PON1-N253G construct for 36 h; or incubated with or without 10 µg/mL recombinant PNGase F for 5 days. Values are relative to PON1-FL and were normalized to GAPDH levels (means ± SD of three biological replicates). $P$ values are indicated as size of the corresponding circle; Student's $t$-test. NS, not significant.
(**L**) Representative confocal images of H1993-GR grown for 5 days in indicated secretomes from 1 µM gefitinib-treated H1993 cells exogenously treated with or without total 8U PNGase F. GR clones were stained for XBP1 (red) and DAPI (nuclei; green). Orientation to the imaging plane (white boxes) were analyzed by line profiles across each cell.

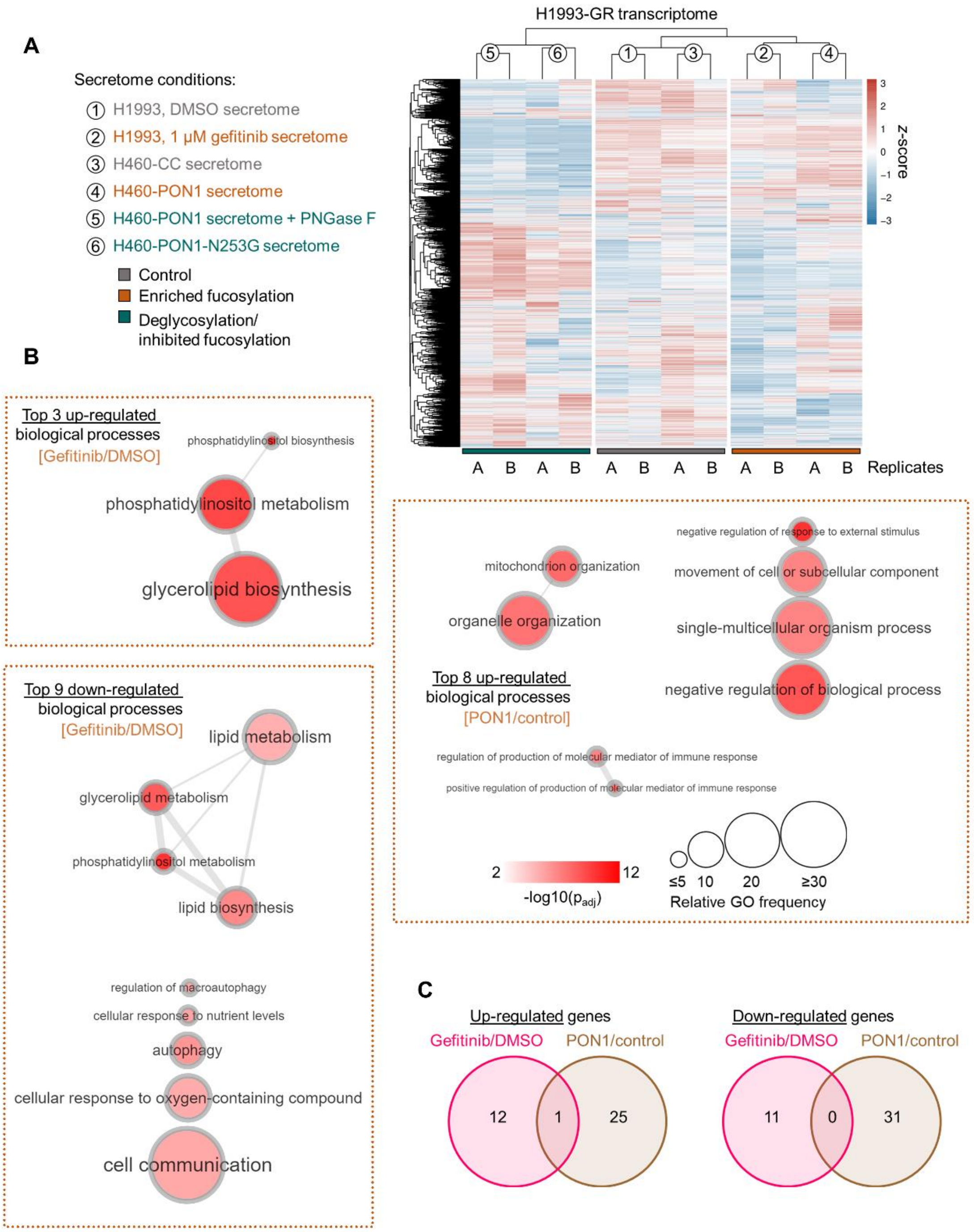


**Supplemental Fig. 22.** RNA-seq of DR clones cultured in PON1 fucosylation-modified secretomes.
(**A**) Heatmap of unsupervised hierarchical clustering from RNA-seq data. Gain and loss signatures correspond to all available genes identified that display indicated z-score.
(**B**) GO analysis of gene expression changes in H1993-GR grown in indicated conditions showing enriched GO terms. Size of circle indicates frequency of the GO term in the underlying GOA database while color indicates adjusted -log10 p-value. Highly similar GO terms are linked by edges in the graph, where the line width indicates the degree of similarity.
(**C**) Venn diagram indicating overlap of up-regulated or down-regulated genes in indicated conditions.

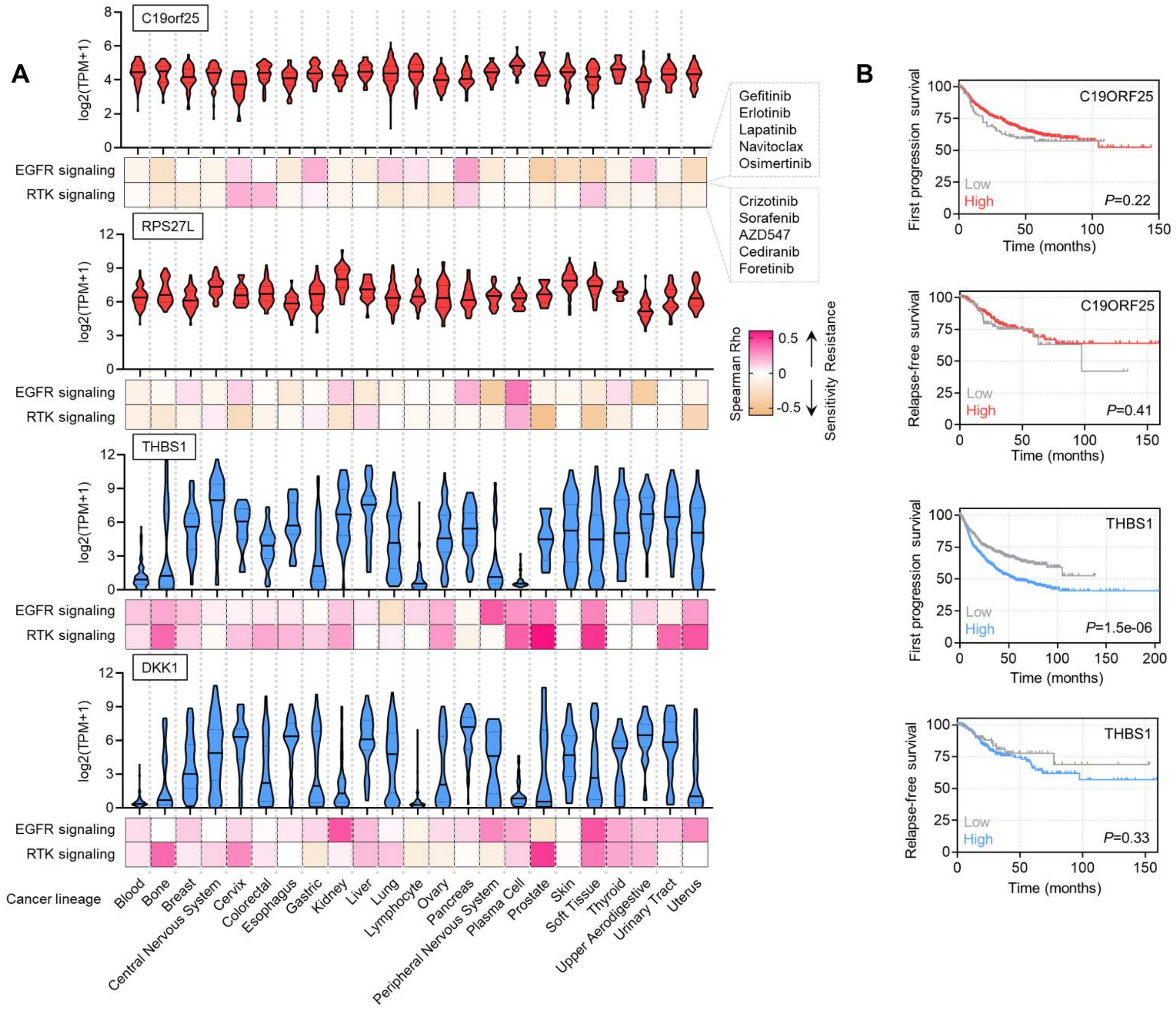


**Supplemental Fig. 23.** Pharmacogenomics data on identified PON1 fucosylation-associated resistance modulator genes.

(**A**) Violin plots depicting pan-cancer expression of indicated top differentially expressed genes from two conditions as in Fig. 5D, in 23 different cancer lineages. Data was obtained from DepMap portal. Bottom shows heatmap of correlation between indicated gene expression and drug response screened in GDSC.

(**B**) Kaplan-Meier plots of FP or RFS in multiple lung cancer patient cohorts. Patient survival data were stratified by indicated gene expression (low or high) in their primary tumors based on microarray (FP) or RNA-seq (RFS) data. P-values were calculated using a log-rank test.

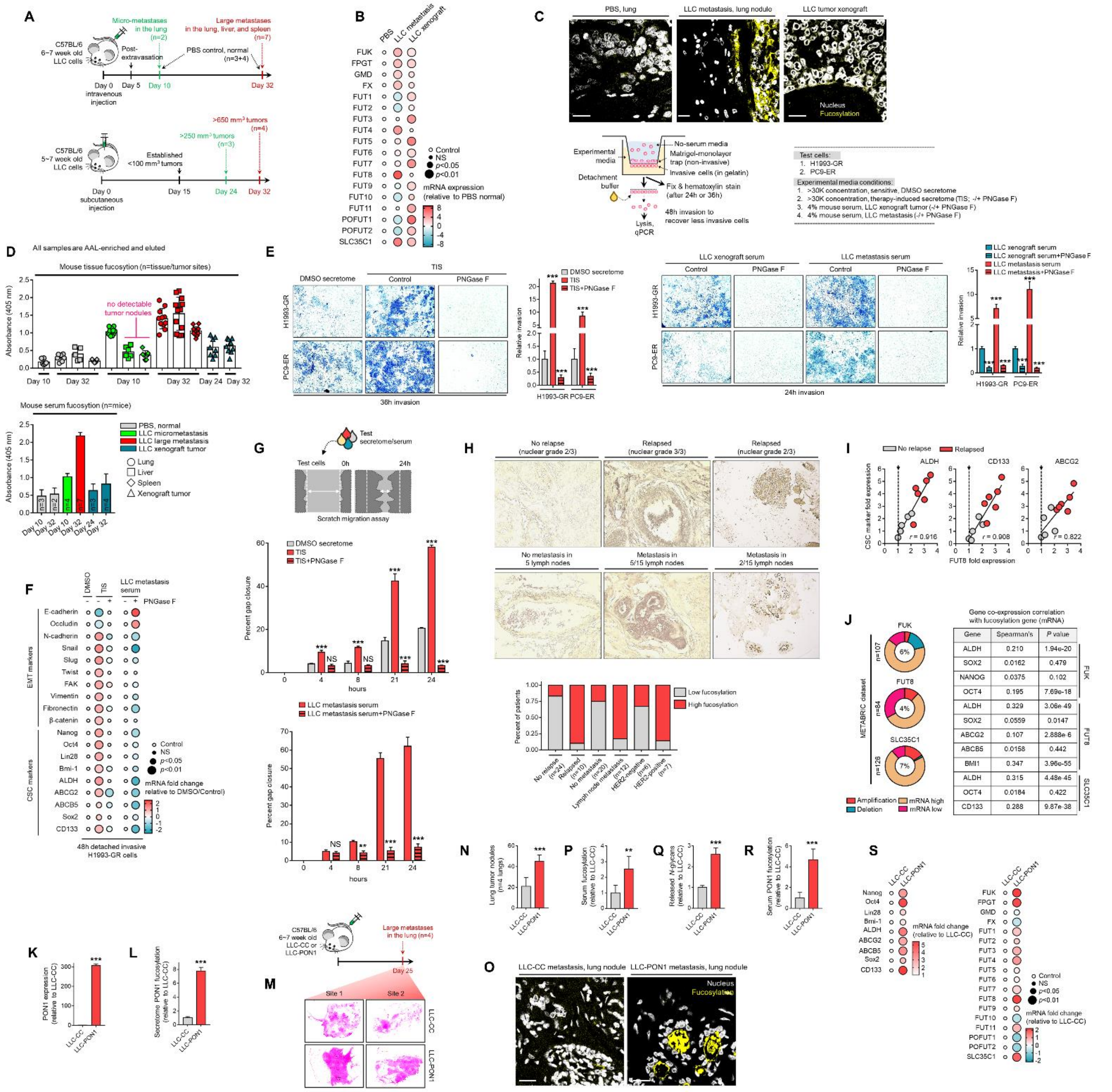


**Supplemental Fig. 24**. Systemic core fucosylation is associated with cancer metastasis and promotes the metastatic potential of DR clones.

(**A**) Schematic of metastatic progression and xenograft induction in C57BL/6 mice using LLC cells.

(**B**) qPCR analysis of indicated fucosylation gene expression in PBS, xenograft tumor, or metastasized tumor tissues. Values indicate mean measurements from three random tumor locations and are relative to PBS and were normalized to GAPDH levels (means ± SD of three biological replicates). *P* values are indicated as size of the corresponding circle; Student's *t*-test. NS, not significant.

(**C**) Representative confocal images of indicated tumor tissue sections stained for core fucosylation (fluorescein-conjugated AAL; yellow) and DAPI (nuclei; white). Scale bars: 100 µm. Representative of three independent experiments.

(**D**) Characterization of fucosylation by sandwich ELLA in tissues or crude sera from indicated mouse models. Values indicate mean absorbance at 405 nm (means ± SD of two biological replicates per sample). Representative of two independent experiments.

(**E**) Transwell Matrigel invasion assay of indicated DR clones prepared as in the schematic. Values are relative to DMSO or xenograft serum (means ± SD of two biological replicates). ****P*<0.001, Student's *t*-test. NS, not significant.

(**F**) qPCR analysis of indicated gene expression in DR clones invaded through the matrix and grown under

indicated conditions as in E. Values are relative to DMSO or non-treated LLC metastasis serum and were normalized to GAPDH levels (means ± SD of three biological replicates). *P* values are indicated as size of the corresponding circle; Student's *t*-test. NS, not significant.
(**G**) Scratch migration assay of H1993-GR in the presence of indicated secretome or serum conditions. Values are relative to DMSO or non-treated LLC metastasis serum (means ± SD of two biological replicates). ***$P<0.001$, Student's *t*-test. NS, not significant.
(**H**) Immunohistochemical analysis of fucosylation in indicated FFPE tumor tissue sections from patients with breast cancer who underwent sequential multidrug therapy. All sections were photographed with an inverted phase-contrast microscope (original magnification, ×200). Scored IHC expression of fucosylation in FFPE tumor sections of relapsed/recurred or non-relapsed breast cancer patients.
(**I**) Co-expression qPCR analysis of indicated CSC genes and FUT8 in the same FFPE tumor tissue sections as in H. Pearson correlation is shown. Representative of two independent experiments. All fold expression values (relapsed/no relapse) are statistically significant ($P<0.01$).
(**J**) OncoPrints visualizing indicated fucosylation gene alterations in the METABRIC dataset and co-occurrence analysis of indicated CSC gene mRNA alterations with indicated fucosylation genes.
(**K**) qPCR analysis of PON1 expression in indicated LLC cells. Values are relative to LLC-CC and were normalized to GAPDH levels (means ± SD of three biological replicates). ***$P<0.001$, Student's *t*-test.
(**L**) HLE analysis of PON1 fucosylation in indicated secretomes derived from LLC cells. Values are relative to LLC-CC (means ± SD of two biological replicates). ***$P<0.001$, Student's *t*-test.
(**M**) Schematic of metastatic progression in C57BL/6 mice using LLC-CC and LLC-PON1 cells and hematoxylin and eosin (H&E) immunohistochemical analysis of large tumor metastasis of indicated LCC cells.
(**N**) Quantification of metastasized tumor nodule formation in indicated mice as in M. Values indicated absolute counts per lung (means ± SD of four mice lungs). ***$P<0.001$, Student's *t*-test.
(**O**) Representative confocal images of indicated tumor tissue sections stained for core fucosylation (fluorescein-conjugated AAL; yellow) and DAPI (nuclei; white). Scale bars: 100 μm.
(**P**) Characterization of fucosylation by sandwich ELLA in crude sera from indicated mouse models as in M. Values are relative to LLC-CC (means ± SD of two biological replicates). **$P<0.01$, Student's *t*-test.
(**Q**) N-glycan release assay in crude sera from indicated mouse models as in M. Values are relative to LLC-CC (means ± SD of three biological replicates). ***$P<0.001$, Student's *t*-test.
(**R**) HLE analysis of PON1 fucosylation in crude sera from indicated mouse models as in M. Values are relative to LLC-CC (means ± SD of three biological replicates). ***$P<0.001$, Student's *t*-test.
(**S**) qPCR analysis of indicated fucosylation gene expression in metastasized tumor tissues from indicated mouse models as in M. Values indicate mean measurements from four random tumor locations and are relative to PBS and were normalized to GAPDH levels (means ± SD of three biological replicates). *P* values are indicated as size of the corresponding circle; Student's *t*-test. NS, not significant.

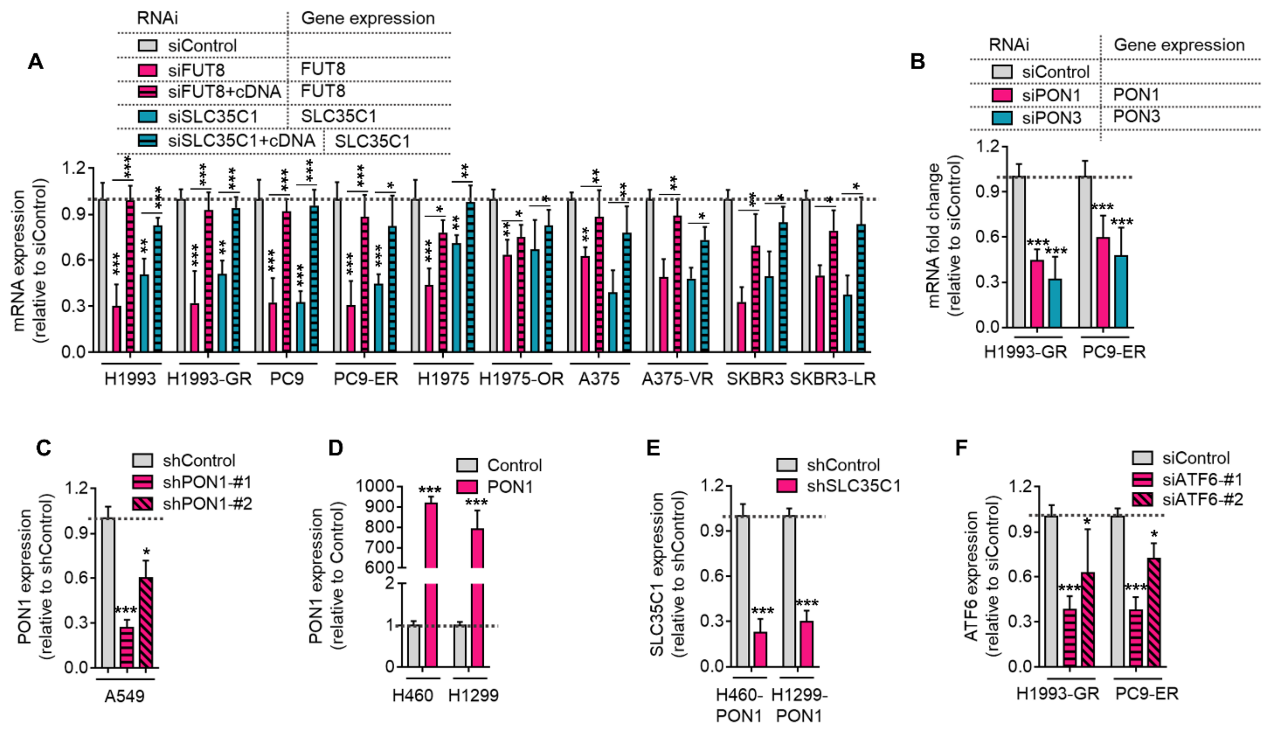


**Supplemental Fig. 25.** Validation of target gene control upon RNAi or stable overexpression in cells.
(**A**) qPCR analysis of FUT8 or SLC35C1 expression in indicated cells or DR clones upon indicated RNAi for 48 h. Values are relative to siControl (means ± SD of two biological replicates). **P*<0.05, ***P*<0.01, ****P*<0.001, Student's *t*-test.
(**B**) qPCR analysis of PON1 or PON3 expression in indicated DR clones upon indicated RNAi for 48 h. Values are relative to siControl (means ± SD of three biological replicates). ****P*<0.001, Student's *t*-test.
(**C**) qPCR analysis of PON1 expression in A549 cells upon stable PON1 knockout. Values are relative to shControl (means ± SD of three biological replicates). **P*<0.05, ****P*<0.001, Student's *t*-test.
(**D**) qPCR analysis of PON1 expression in H460 or H1299 cells upon stable PON1 overexpression. Values are relative to control (means ± SD of three biological replicates). ****P*<0.001, Student's *t*-test.
(**E**) qPCR analysis of PON1 expression in PON1-overexpressing H460 or H1299 cells upon stable SLC35C1 knockout. Values are relative to shControl (means ± SD of two biological replicates). ****P*<0.001, Student's *t*-test.
(**F**) qPCR analysis of ATF6 expression in indicated DR clones upon ATF6 RNAi for 48 h. Values are relative to siControl (means ± SD of three biological replicates). **P*<0.05, ****P*<0.001, Student's *t*-test.

**Supplemental Movie 1.** 3D spheroid formation of CellTracker-Green-labeled H1993 and CellTracker-Red-labeled H1993-GR admixture within 24 h.

**Supplemental Movie 2.** Live-imaging of CellTracker-Green-labeled H1993 and CellTracker-Red-labeled H1993-GR admixture upon treatment with 2 μM gefitinib within 24 h.

**Supplemental Movie 3.** Live-imaging of CellTracker-Green-labeled PC9 and CellTracker-Red-labeled PC9-ER admixture upon treatment with 0.1 μM erlotinib within 24 h.
