## Supplemental Tables for "Widespread multi-targeted therapy resistance via drug-induced secretome fucosylation"

**Table S1. Information of human lung cancer patients (cohort for sera and plasma).**

| Study ID | Age | Sex | Blood draw date | Treatment |
|---|---|---|---|---|
| 1 | 83 | M | 7/9/2012 | - |
| 2 | 72 | M | 7/12/2012 | - |
| 3 | 57 | F | 5/23/2013 | - |
| 4 | 73 | F | 3/27/2013 | - |
| 5 | 61 | M | 10/17/2013 | - |
| 6 | 59 | M | 8/27/2018 | - |
| 7 | 72 | M | 1/3/2013 | - |
| 8 | 52 | F | 4/9/2013 | - |
| 9 | 59 | M | 6/29/2016 | - |
| 10 | 66 | F | 8/18/2014 | - |
| 23 | 36 | F | 3/18/2014 | - |
| 24 | 48 | F | 2015-0121 | - |
| 25 | 81 | F | 8/14/2013 | - |
| 26 | 45 | F | 5/22/2013 | - |
| 27 | 37 | F | 8/22/2013 | - |
| 28 | 72 | F | 8/8/2014 | - |
| 29 | 60 | F | 4/8/2013 | - |
| 30 | 71 | M | 6/19/2013 | - |
| 31 | 37 | F | 9/1/2015 | - |
| 32 | 59 | M | 10/7/2014 | - |
| 33 | 64 | F | 9/2/2014 | - |
| 34 | 48 | M | 7/18/2016 | - |
| 35 | 59 | F | 6/24/2014 | - |
| 36 | 52 | M | 9/4/2018 | - |
| 37 | 73 | M | 7/28/2016 | - |
| 38 | 57 | M | 10/20/2017 | - |
| 39 | 75 | F | 8/3/2018 | - |
| 11 | 63 | F | 20180914 | Osimertinib |
| 12 | 63 | F | 20181112 | Osimertinib |
| 13 | 63 | F | 20190201 | Osimertinib |
| 14 | 61 | F | 20180713 | Osimertinib |
| 15 | 61 | F | 20180914 | Osimertinib |
| 16 | 42 | F | 20180302 | Osimertinib |
| 17 | 42 | F | 20180430 | Osimertinib |
| 18 | 50 | F | 20180124 | Osimertinib |

| Study ID | Age | Sex | Blood draw date | Treatment |
|---|---|---|---|---|
| 19 | 50 | F | 20180326 | Osimertinib |
| 20 | 69 | F | 20180115 | Osimertinib |
| 21 | 69 | F | 20180312 | Osimertinib |
| 22 | 45 | F | 20180413 | Osimertinib |
| 40 | 45 | F | 20180612 | Osimertinib |
| 41 | 45 | F | 20180720 | Osimertinib |
| 42 | 58 | M | 20180309 | Osimertinib |
| 43 | 58 | M | 20180504 | Osimertinib |
| 44 | 70 | M | 20180327 | Osimertinib |
| 45 | 70 | M | 20180524 | Osimertinib |
| 46 | 71 | M | 20180516 | Osimertinib |
| 47 | 71 | M | 20180720 | Osimertinib |
| 48 | 71 | M | 20181112 | Osimertinib |
| 49 | UNKNOWN | UNKNOWN | 20180611 | Osimertinib |
| 50 | UNKNOWN | UNKNOWN | 20180806 | Osimertinib |
| 51 | UNKNOWN | UNKNOWN | 20180727 | Osimertinib |
| 52 | UNKNOWN | UNKNOWN | 20180921 | Osimertinib |
| 53 | UNKNOWN | UNKNOWN | 20181123 | Osimertinib |
| 54 | UNKNOWN | UNKNOWN | 20180827 | Osimertinib |
| 55 | UNKNOWN | UNKNOWN | 20181017 | Osimertinib |
| 56 | 76 | M | 20190213 | Osimertinib |
| 57 | 76 | M | 20190410 | Osimertinib |

**Table S2. Clinicopathologic information of human lung cancer patients (cohort for tumor tissues).**

| Study ID | Age | Sex | Smoking | Type | T/N/M | Stage | Relapse | Relapse-free period (months) |
|---|---|---|---|---|---|---|---|---|
| 800 | 22 | M | Current | A | UNKNOWN | 2 | X | 26 |
| 817 | 63 | F | Never | A | UNKNOWN | 1 | X | 26 |
| 826 | 52 | F | Never | A | 1/0/0 | 1 | X | 26 |
| 982 | 70 | F | Former | A | 4/0/0 | 1 | X | 22 |
| 862 | 70 | M | Current | S | 3/1/0 | 2 | X | 26 |
| 923 | 51 | F | Never | A | UNKNOWN | 3 | X | 23 |
| 927 | 84 | M | Former | S | 3/0/0 | 1 | X | 23 |
| 932 | 69 | M | Former | S | 2/1/0 | 2 | X | 23 |
| 936 | 56 | M | Never | A | 3/1/0 | 2 | X | 23 |
| 940 | 69 | M | Former | A | UNKNOWN | 2 | X | 23 |
| 857 | 75 | M | Former | A | 3/0/0 | 1 | X | 25 |
| 877 | 76 | F | Never | A | UNKNOWN | 1 | X | 25 |
| 912 | 60 | M | Never | A | UNKNOWN | 1 | X | 25 |
| 953 | 53 | F | Never | A | 2/0/0 | 1 | X | 22 |
| 956 | 35 | F | Never | L | UNKNOWN | 2 | X | 22 |
| 959 | 55 | M | Former | A | 3/0/0 | 1 | X | 22 |
| 969 | 76 | M | Current | A | 3/0/0 | 1 | X | 22 |
| 971 | 69 | M | Former | S | 2/0/0 | 1 | X | 22 |
| 941 | 68 | M | Former | A | 2/1/0 | 2 | X | 23 |
| 942 | 51 | M | Current | A | UNKNOWN | 1 | X | 23 |
| 992 | 84 | M | Former | S | 4/0/0 | 2 | X | 20 |
| 993 | 38 | M | Current | A | UNKNOWN | 3 | X | 20 |
| 996 | 64 | M | Never | A | UNKNOWN | 3 | X | 20 |
| 999 | 66 | F | Never | A | 2/0/0 | 1 | X | 20 |
| 1003 | 72 | M | Current | S | UNKNOWN | 3 | X | 20 |
| 1009 | 78 | F | Never | A | 5/0/0 | 2 | X | 20 |
| 815 | 48 | M | Current | A | 5/2/0 | 3 | X | 23 |
| 866 | 77 | M | Former | A | 1/0/0 | 1 | X | 23 |
| 851 | 65 | M | Former | S | 3/0/0 | 1 | X | 23 |
| 889 | 67 | M | Never | A | 3/0/0 | 1 | X | 23 |
| 899 | 72 | M | Former | S | 2/0/0 | 1 | X | 23 |
| 831 | 71 | M | Current | S | 3/0/0 | 1 | X | 26 |
| 915 | 57 | M | Current | S | 1/0/0 | 1 | X | 24 |
| 894 | 70 | F | Never | A | 2/0/0 | 1 | X | 22 |
| 888 | 68 | F | Never | A | 3/0/0 | 1 | X | 22 |
| 983 | 60 | M | Former | A | 1/0/0 | 1 | X | 22 |
| 986 | 67 | M | Former | A | 3/0/0 | 1 | X | 22 |
| 880 | 56 | M | Former | A | 2/0/0 | 1 | O | 13 |
| 870 | 72 | M | Former | A | 1/0/0 | 1 | O | 19 |
| 943 | 66 | M | Never | S | 3/0/0 | 1 | O | 7 |
| 794 | 75 | M | Never | A | 5/2/0 | 3 | O | 8 |
| 881 | 67 | F | Never | A | 3/0/0 | 1 | O | 15 |
| 882 | 45 | F | Never | A | 3/1/0 | 2 | O | 13 |
| 937 | 64 | M | Current | S | 2/0/0 | 1 | O | 13 |
| 887 | 76 | M | Former | S | 4/0/0 | 2 | O | 18 |
| 863 | 43 | F | Never | A | 3/2/1 | 3 | O | 13 |
| 839 | 45 | M | Current | A | 3/0/0 | 3 | O | 7 |
| 836 | 82 | M | Never | A | 2/1/0 | 2 | O | 13 |
| 984 | 71 | F | Former | A | 1/0/0 | 1 | O | 18 |
| 1000 | 56 | F | Never | A | 3/2/0 | 3 | O | 6 |
| 1007 | 65 | M | Former | A | 1/0/0 | 1 | O | 14 |
| 917 | 62 | M | Current | A | 4/0/0 | 2 | O | 19 |
| 828 | 80 | M | Current | S | 2/0/0 | 1 | X | 26 |

### Table S3 Clinicopathologic information of human breast cancer patients (cohort for tumor tissues).

| Study ID | Age | Nuclear grade | Histologic grade | Tumor size | Lymph node metastasis | Recurrence/relapse |
|---|---|---|---|---|---|---|
| 12 | 36 | 3/3 | III/III | 5.1 x 5.0 x 3.0cm, 2.2 x 1.4 x 1.5cm | metastasis in one out of 28 lymph nodes | O |
| 1 | 41 | 3/3 | III/III | 3.8 x 3.5 x 4.5cm | metastasis in one out of 10 lymph nodes | N/A |
| 2 | 44 | 3/3 | II/III | up to 2.3 x 1.1cm | metastasis in two out of 16 lymph nodes | X |
| 11 | 31 | 3/3 | II/III | 1.8 x 1.2 x 3.0cm | metastasis in one out of 22 lymph nodes | O |
| 3 | 48 | 3/3 | II/III | up to 2.1 x 1.4 x 2.0cm | metastasis in ten out of 24 lymph nodes with extracapsular extension | X |
| 4 | 65 | 3/3 | II/III | 4.1 x 1.7 x 2.0cm | metastasis in three out of 19 lymph nodes | X |
| 10 | 66 | 3/3 | III/III | 4.8 x 3.6 x 3.5cm | no metastasis in 19 lymph nodes | O |
| 9 | 43 | 3/3 | III/III | 4.5 x 2.7 x 4.0cm | no metastasis in 5 lymph nodes | O |
| 5 | 62 | 3/3 | III/III | up to 1.9 x 1.6 x 2.5cm | metastasis in one out of nine lymph nodes | X |
| 6 | 45 | 3/3 | III/III | up to 4.8 x 3.8 x 3.0cm | no metastasis in five lymph nodes | N/A |
| 31 | UNKNOWN | UNKNOWN | UNKNOWN | UNKNOWN | UNKNOWN | O |
| 7 | 47 | 3/3 | III/III | up to 0.3 x 0.3 x 0.5cm | metastasis in five out of 15 lymph nodes | O |
| 13 | 57 | 3/3 | III/III | 0.3 x 0.2cm | no metastasis in five lymph nodes | X |
| 33 | 26 | 3/3 | III/III | up to 0.2 x 0.2 x 0.2cm | no metastasis in nine lymph nodes | O |
| 14 | 42 | 2/3 | II/III | 0.2 x 0.2 x 0.2cm and 0.1 x 0.1 x 0.1cm | no metastasis in seven lymph nodes | X |
| 15 | 58 | 2/3 | III/III | 0.3 x 0.2 x 0.2cm | no metastasis in nine lymph nodes | X |
| 16 | 40 | 2/3 | II/III | up to 0.4 x 0.3cm | metastasis in one out of 14 lymph nodes | X |
| 32 | 63 | 2/3 | II/III | up to 0.3 x 0.2cm | no metastasis in 11 lymph nodes | O |
| 17 | 56 | 3/3 | III/III | 0.4 x 0.3 x x0.5 cm | 0/3 | X |
| 18 | 51 | 2/3 | II/III | up to 0.4 x 0.4 cm | 3 with extracapsular extension/13 | X |
| 19 | 53 | 3/3 | III/III | 0.4 x 0.4 x 0.4cm | no metastasis in eight lymph nodes | X |
| 20 | 48 | 3/3 | II/III | up to 0.2 x 0.2 x 0.2 cm | 4/14 | X |
| 21 | 37 | 2/3 | II/III | 0.3 x 0.3cm | 0/3 | X |
| 22 | 42 | 2/3 | II/III | 0.3 x 0.2cm | no metastasis in 21 lymph nodes | X |
| 23 | 38 | 2/3 | II/III | up to 0.3 x 0.2cm | metastasis in 11 out of 20 lymph nodes | X |
| 24 | 31 | 3/3 | III/III | up to 0.2 x 0.2 cm | 0/5 | X |
| 25 | 38 | 2/3 | II/III | 0.2 x 0.2 cm | 0/4 | X |
| 26 | 59 | 2/3 | II/III | up to 0.4 x 0.3cm | no metastasis in six lymph nodes | X |
| 8 | 48 | 3/3 | III/III | 0.2 x 0.2 cm | 0/5 | O |
| 27 | 31 | 3/3 | II/III | up to 0.4 x 0.2 cm | 5/12 | X |
| 28 | 38 | 3/3 | III/III | 0.4 x 0.3cm | no metastasis in three lymph nodes | X |
| 29 | 54 | 2/3 | II/III | 0.4 x 0.3 cm | 0/8 | X |
| 30 | 47 | 3/3 | III/III | 0.4 x 0.4 x 0.4cm | no metastasis in nine lymph nodes | X |

**Table S4.** siRNA and shRNA sequences.

| Target gene | RNAi | Sequence (5′→3′) | # |
|---|---|---|---|
| FUT8 | siRNA | UGG AGC UAA AGA GCU CUG GTT | 1 |
| FUT8 | siRNA | UCU CAG AAU UGG CGC UAU GTT | 2 |
| SLC35C1 | siRNA | GUC CCU AAC AUC ACA UUG U | 1 |
| SLC35C1 | siRNA | GCC UGA CUU UCU ACA ACA A | 2 |
| PON1 | siRNA | GCU CUG GAU UAA AGU AUC CUG GAA U | 1 |
| PON1 | siRNA | AAG ACU GGU GGU UCC UGA AGA GUG C | 2 |
| PON3 | siRNA | GUG GCU CUG AAG AUA UUG A | 1 |
| PON3 | siRNA | CAC CGU GUA UGC CAA CAA U | 2 |
| ATF6 | siRNA | GCU CUC UUU GUU GUU GCU UAG UGG A | 1 |
| ATF6 | siRNA | CAG AGU CUC UCA GGU UAA A | 2 |
| PON1 | shRNA | CCT AAA GTG ACA CAG GTT TAT | 1 |
| PON1 | shRNA | GCT TTC ATT AGC TCT GGA TTA | 2 |
| SLC35C1 | shRNA | CCT GAC TTT CTA CAA CAA CGT | 1 |
| SLC35C1 | shRNA | CGG CCT GTT TGG CTT TGC CAT | 2 |
| Control | siRNA | UUC UCC GAA CGU GUC ACG U | 1 |
| Control | siRNA | AGG UAG UGU AAU CGC CUU G | 2 |
| Control | shRNA | TAG ATA AGC ATT ATA ATT CCT A | 1 |

**Table S5.** Oligonucleotide qPCR primers.

| Target gene | Species | Forward (5′→3′) | Reverse (5′→3′) |
|---|---|---|---|
| FUK | H. sapiens | GGC GAA TCC TAC ACG ACT GAC | GAG TTC TAG CGA GGT TGC TGT GA |
| FPGT | H. sapiens | CAG AGC TCG GCT TAC AGT CC | GCA GTT TTC CCC AAC TGA AA |
| GMD | H. sapiens | CGA GAA GGG CTA CCA GGT TC | GAG ACC ACA GGT ACG GAT GG |
| TSTA3 (FX) | H. sapiens | ACA ATG AAG TGG AGC CCA TC | CAG GTA GGT CCT CAG CTT GC |
| FUT1 | H. sapiens | AAA GCG GAC TGT GGA TCT | GGA CAC AGG ATC GAC AGG |
| FUT2 | H. sapiens | ATC ATG ACC ATT GGG ACG TT | GTG CTT GAG TAA GGG GGA CA |
| FUT3 | H. sapiens | GCT CAG AGT TCA GAC AGG TCC AA | CCA GAA GTC CTC ATT TAC AGT CGA T |
| FUT4 | H. sapiens | TCC TAC GGA GAG GCT CAG | TCC TCG TAG TCC AAC ACG |
| FUT5 | H. sapiens | GGT GTC ACA AAT GCA TCA CTA TGG | ACA TCC ACA GGC ACT ATC AGA GT |
| FUT6 | H. sapiens | CAT TTC TGC TGC CTC AGG | GGG CAA GTC AGG CAA CTC |
| FUT7 | H. sapiens | CCA CGA TCA CCA TCC TTG | AGG CTT CGG TTG GCA CTC |
| FUT8 | H. sapiens | TCT AGC CGA GAA CTG TCC | GCT GCT CTT CTA AAA CGC |
| FUT9 | H. sapiens | CAA CTG TCT ACG TGC TTC | AGA CAT GCC ATG AAA CAG |
| FUT10 | H. sapiens | ATC TTG CCT GTG CGT CAC | ATG CGT AGG TGC TTC CTC |
| FUT11 | H. sapiens | CTC TTG GCT TTC TTG TCC | ATG ACG GAG TGA TTG TTC |
| POFUT1 | H. sapiens | AAC CAG GCC GAT CAC TTC TTG | GTT GGT GAA AGG AGG CTT GTG |
| POFUT2 | H. sapiens | AAA GAA GGG ACC TGG GAA GA | GGC TGA TGT GTT TCT CAG CA |
| SLC35C1 | H. sapiens | AAG ATC AAG AGG CCA GCA GA | GGA AGG AGC TTG CTT GTT TG |
| P16INK4A | H. sapiens | AAT CTC CGC GAG GAA AGC | GTC TGC AGC GGA CTC CAT |
| LMNB1 | H. sapiens | GGG AAG TTT ATT CGC TTG AAG A | ATC TCC CAG CCT CCC ATT |
| IL1α | H. sapiens | TCC ATA ACC CAT GAT CTG GAA | TTG GTT GAG GGA ATC ATT CAT |
| IL6 | H. sapiens | GCT ACC AAA CTG GAT ATA ATC AGG A | CCA GGT AGC TAT GGT ACT CCA GAA |
| MMP3 | H. sapiens | CAA AAC ATA TTT CTT TGT AGA GGA CAA | TTC AGC TAT TTG CTT GGG AAA |
| MMP9 | H. sapiens | ACG ACA TAG ACG GCA TCC A | GCT GTG GTT CAG TTG TGG TG |
| CXCL1 | H. sapiens | AGG GAA TTC ACC CCA AGA AC | TGG ATT TGT CAC TGT TCA GCA |
| CXCL10 | H. sapiens | GCT GCC GTC ATT TTC TGC | TCT CAC TGG CCC GTC ATC |
| IGF1 | H. sapiens | GGC ATA GCT GGC CAA ACA A | CAC TTG GGA GAA GGC TTA GAA TAA A |
| BMP2 | H. sapiens | ATG GAT TCG TGG TGG AAG TG | GTG GAG TTC AGA TGA TCA GC |
| MTPN | H. sapiens | GCC AAG GGA GAA GAT GTC AA | AGC ACC CTT TGA CAG AAG CA |
| ESR1 | H. sapiens | CCA TGA CCC TCC ACA CC | CTC GTT CCC TTG GAT CTG A |
| NFKBIA | H. sapiens | GAG TTA CCT ACC AGG GCT ATT C | CTC TCC TCA TCC TCA CTC TCT |
| PRL | H. sapiens | ACA ATG GAG CTT CAT GTC CTC | TCC ATG AAT AGG AGA GTT CTT TAG TTT |
| AGT | H. sapiens | TTG TTG AGA GCT TGG GTC CCT TCA | CAG ACA CTG AGG TGC TGT TGT CCA |
| MAPK14 | H. sapiens | GCA GGG ACC TTC TCA TAG AT | GAG GGA TAG CCT CAG ACC |
| TP73 | H. sapiens | GGA AGA TGG CCC AGT CCA CCG | GTG GAT CTC GGC CTC CGT GAA C |
| GHI | H. sapiens | CCT AGA GGA AGG CAT CCA AA | GCA GCC CGT AGT TCT TGA GTA G |
| ATF4 | H. sapiens | AAA CCT CAT GGG TTC TCC AG | GGC ATG GTT TCC AGG TCA TC |
| SHH | H. sapiens | GAA AGC AGA GAA CTC GGT GG | GGA AAG TGA GGA AGT CGC TG |
| PON1 | H. sapiens | GGA TCC ATG GCG AAG CTG ATT GCG TCT AC | GCG GCC GCG AGC TCA CAG TAA AGA GCT TTG |
| PON3 | H. sapiens | CCA GGC ATG CCA AAC TTT GC | GGG CCA AAA ATA CTA GTG TTC |
| ATF6 | H. sapiens | AAT TCT CAG CTG ATG GCT GT | TGG AGG ATC CTG GTG TCC AT |
| ATF2 | H. sapiens | TGC CTG TTG CTA TTC CTG C | GCT CTT CTC CGA CGA CCA CT |
| ATF3 | H. sapiens | ATG ATG CTT CAA CAC CCA GG | TTT CGG CAC TTT GCA GCT G |
| AR | H. sapiens | TAC CAG CTC ACC AAG CTC CTG | GAA AGT CCA CGC TCA CCA TGT G |
| E2F | H. sapiens | GGA TTT CAC ACC TTT TCC TGG AT | CCT GGA AAC TGA CCA TCA GTA CCT |
| HIF | H. sapiens | CAT AAA GTC TGC AAC ATG GAA GGT | ATT TGA TGG GTG AGG AAT GGG TT |
| STAT3 | H. sapiens | CAT GTC AAA GGT GAG GGA CTC | CCC CGC ACT TTA GAT TCA TT |
| ISRE | H. sapiens | GAG ATG AAT GAG TGC TGT TTT GG | AGA AGG GGC AGG GAA GTA ATG |
| LXR | H. sapiens | AAG CCC TGC ATG CCT ACG T | TGC AGA CGC AGT GCA AAC A |
| PR | H. sapiens | GGC GAG AGG CAA CTT CTT TC | CAT CTG CCC ACT GAC GTG TT |

| | | | |
|---|---|---|---|
| PPAR | H. sapiens | TCG GCG AAC TAT TCG GCT G | GCA CTT GTG AAA ACG GCA GT |
| PERK | H. sapiens | AAA AAG CAG TGG GAT TTG GA | CTG GAA TAT ACC GAA GTT CAA AG |
| CHOP | H. sapiens | GCT GTG GTA GTG AGC TGT TGC A | CAC AGC CCA GGT ATG GAA TCA |
| GRP94 | H. sapiens | ACT GTT GAG GAG CCC ATG GAG G | GCT GAA GAG TCT CGC GGG AAA C |
| GRP78 | H. sapiens | GTG TTC AAG AAC GGC CGC GTG | GTT TGC CCA CCT CCA ATA TCA AC |
| HEDJ | H. sapiens | GAG TCA CTG GTT GGC TTT GA | CCT TTC TTC CAT AGC TTC GC |
| P58IPK | H. sapiens | GAA GCA TCT TGA ATT GGG GA | CAA GCT TCC CTT GTT TGA GC |
| ERDJ4 | H. sapiens | AAA ATA AGA GCC CGG ATG CT | CAG TCC TGC AGT GCT TGC TA |
| XBP1 | H. sapiens | TGA AAA ACA GAG TAG CAG CTC AGA | CCC AAG CGC TGT CTT AAC TC |
| CDH1 (E-cadherin) | H. sapiens | GAA CAG CAC GTA CAC AGC CCT | GCA GAA CTG TCC CTG TCC CAG |
| CDH2 (N-cadherin) | H. sapiens | GAC GGT TCG CCA TCC AGA C | TCG ATT GGT TTG ACC ACG G |
| OCLN (Occludin) | H. sapiens | CCC ATC TGA CTA TGT GGA AAG AG | AAC CGG CGT GGA TTT ATA GG |
| SNAI1(Snail) | H. sapiens | CAT GTC TGG ACC TGG TTC CT | AAG GGT CCT TGA GGG AGG TA |
| SNAI2 (Slug) | H. sapiens | TGG TTG CTT CAA GGA CAC AT | GTT GCA GTG AGG GCA AGA A |
| TWST1 (Twist) | H. sapiens | GTC CGC AGT CTT ACG AGG AG | GCT TGA GGG TCT GAA TCT TGC T |
| FAK | H. sapiens | GTG CTC TTG GTT CAA GCT GGA T | ACT TGA GTG AAG TCA GCA AGA TG |
| VIM (Vimentin) | H. sapiens | TAC AGG AAG CTG CTG GAA GG | ACC AGA GGG AGT GAA TCC AG |
| FN1 (Fibronectin) | H. sapiens | GGA AAG TGT CCC TAT CTC TGA TAC C | AAT GTT GGT GAA TCG CAG GT |
| CTNNB1 (β-catenin) | H. sapiens | GTG CAA TTC CTG AGC TGA CA | CTT AAA GAT GGC CAG CAA GC |
| NANOG | H. sapiens | CCT CCT CCA TGG ATC TGC TTA TTC A | CAG GTC TTC ACC TGT TTG TAG |
| OCT4 | H. sapiens | ACA TCA AAG CTC TGC AGA AAG AAC T | CTG AAT ACC TTC CCA AAT AGA ACC C |
| LIN28 | H. sapiens | TTG TCT TCT ACC CTG CCC TCT | GAA CAA GGG ATG GAG GGT TTT |
| BMI1 | H. sapiens | CCA GGG CTT TTC AAA AAT GA | CCG ATC CAA TCT GTT CTG GT |
| ALDH | H. sapiens | TGT TAG CTG ATG CCG ACT TG | TTC TTA GCC CGC TCA ACA CT |
| ABCG2 | H. sapiens | CAG GTC TGT TGG TCA ATC TCA CA | TCC ATA TCG TGG AAT GCT GAA G |
| ABCB5 | H. sapiens | TGG ATC CAA CAC CTC TAT GCT AAA | GGC AGG TTT TCT CGA TGA ACT G |
| SOX2 | H. sapiens | AGC TAC AGC ATG ATG CAG GA | GGT CAT GGA GTT GTA CTG CA |
| CD133 | H. sapiens | TGG ATG CAG AAC TTG ACA ACG T | ATA CCT GCT ACG ACA GTC GTG GT |
| GAPDH | H. sapiens | CTC TCT GCT CCT CCT GTT CGA C | TGA GCG ATG TGG CTC GGC T |
| TUBB | H. sapiens | GCC TTC CTT CTT GGG TAT GG | GCA CTG TGT TGG CAT AGA GG |
| 18S | H. sapiens | CGG CGA CGA CCC ATT CGA AC | GAA TCG AAC CCT GAT TCC CCG TC |
| Pon1 | M. musculus | GCA TCT GAA AAC CAT CAC ACA | AAG CTC TCA GGT CCA ATA GCA |
| Nanog | M. musculus | AGG GTC TGC TAC TGA GAT GCT CTG | CAA CCA CTG GTT TTT CTG CCA CCG |
| Oct4 | M. musculus | TGG AGA CTT TGC AGC CTG AG | TGA ATG CAT GGG AGA GCC CA |
| Lin28 | M. musculus | GTT CGG CTT CCT GTC TAT GA | GTT GTA GCA CCT GTC TCC TT |
| Bmi1 | M. musculus | TTT TAT GCT GAA CGA CTT TTA ACT T | GCT CAG TGA TCT TGA TTC TGG T |
| Aldh | M. musculus | GAC AGG CTT TCC AGA TTG GCT C | AAG ACT TTC CCA CCA TTG AGT GC |
| Abcg2 | M. musculus | TCG CAG AAG GAG ATG TGT TGA G | CCA GAA TAG CAT TAA GGC CAG G |
| Abcb5 | M. musculus | GTG GCT GAA GAA GCC TTG TC | TGA AGC CGT AGC CCT CTT TA |
| Sox2 | M. musculus | AAA GCG TTA ATT TGG ATG GG | ACA AGA GAA TTG GGA GGG GT |
| Cd133 | M. musculus | TTG GTG CAA ATG TGG AAA AG | ATT GCC ATT GTT CCT TGA GC |
| Fuk (fcsk) | M. musculus | GGA CCA AGT CAG TGG CCT AA | GAG CGT ACC AGT TCC TCA GC |
| Fpgt | M. musculus | TTG TTG GAC ATG AGG CAG AA | TAA GCC CAG CTC CGT CTT TA |
| Gmd | M. musculus | GGA GCA GCC AAA CTC TAT GC | TTC CCA AGC TGA AAC ATT CC |
| Tsta3 (fx) | M. musculus | GGG TGT TTG TCT CCT CCA AA | GGA CGT TGT CAT TGA TGT GC |
| Fut1 | M. musculus | GCA TCC GCC CTC ATA CCT | GCC AGC GAA GAC CAC ATC A |
| Fut2 | M. musculus | CCC ACT TCC TCA TCT TTG TCT TT | TTT GAA CCG CCT GTA ATT CCT T |
| Fut3 (sec1) | M. musculus | GGT TTT CAA GCC AGA AGC AG | AGA GCA CCA GGA AGG CAG TA |
| Fut4 | M. musculus | CAG CCT GCG CTT CAA CAT C | CGC CTT ATC CGT GCG TTC T |
| Fut7 | M. musculus | GTA GGG GCC CAG GAT AAG AG | CCT TGA GGC TAA GCA CCT TG |
| Fut8 | M. musculus | CAG GGG ATT GGC GTG AAA AAG | CGT GAT GGA GTT GAC AAC CAT AG |
| Fut9 | M. musculus | ATC CAA GTG CCT TAT GGC TTC T | TGC TCA GGG TTC CAG TTA CTC A |
| Fut11 | M. musculus | TAA CTT GGA AGA CTG CGT TAC TG | GGC TGA GAT ACT AGC TCC ATA CC |

| | | | |
|---|---|---|---|
| Pofut1 | M. musculus | TGT ACT CGC AGG ACA AGC ATG AGT | AAC ATG ATC TCG AGT CAG TAC GCA ATC TTC CAG TGT GTG |
| Pofut2 | M. musculus | TGT ACT CGC AGG ACA AGC ATG AGT | AAC ATG ATC TCG AGT CAG TAC GCA ATC TTC CAG TGT GTG |
| Slc35c1 | M. musculus | CCA ACA CCT GAC CCA GCT AT | GAC AGG AGC CTG ATG GGA TA |
| Gapdh | M. musculus | TGT GTC CGT CGT GGA TCT GA | TTG CTG TTG AAG TCG CAG GAG |
| Actb | M. musculus | CAA CGG CTC CGG CAT GTG C | CTC TTG CTC TGG GCC TCG |
